# Single-molecule live cell imaging of Rep reveals the dynamic interplay between an accessory replicative helicase and the replisome

**DOI:** 10.1101/430371

**Authors:** Aisha Syeda, Adam J. M. Wollman, Alex Hargreaves, Janny G. Brüning, Peter McGlynn, Mark C. Leake

**Affiliations:** Department of Biology, University of York, York YO10 5DD, United Kingdom.; Department of Physics, University of York, York YO10 5DD, United Kingdom.

**Author notes:** These authors contributed equally (Aisha Syeda, Adam J. M. Wollman). Contact corresponding author for proofs: Prof Mark Leake, Department of Physics and Biology, University of York, York YO10 5DD, United Kingdom. Orcid ID: http://orcid.org/0000-0002-1715-1249.

**Keywords:** Single-molecule, DNA replication, DNA repair, fluorescent protein, super-resolution, helicase, nucleoprotein barriers, replication restart

## Abstract

DNA replication requires strategies to cope with nucleoprotein barriers that impair the efficient translocation of the replisome. Biochemical and genetic studies indicate accessory helicases play essential roles in continuity of replication in the presence of nucleoprotein barriers, but how they operate in the native cellular environment is unclear. With high-speed single-molecule microscopy we determine the dynamic patterns of localization of genomically-encoded fluorescent protein constructs of the bacterial accessory helicase Rep and core replisome protein DnaQ in live *E. coli* cells. We demonstrate that Rep colocalizes with 70% of replication forks. Colocalisation is dependent upon interaction with replicative helicase DnaB, with an underlying hexameric stoichiometry of Rep indicating maximal occupancy of the single DnaB hexamer within the replisome. We find that Rep associates dynamically with the replisome with an average dwell time of 6.5 ms dependent on ATP hydrolysis, indicating rapid binding then translocation away from the fork. We also imaged the PriC replication restart factor given the known Rep-PriC functional interaction and observe Rep-replisome association is also dependent on the presence of PriC. Our findings suggest two Rep-replisome populations *in vivo:* one involving Rep continually associating with DnaB then translocating away to aid nucleoprotein barrier removal ahead of the fork, another assisting PriC-dependent reloading of DnaB if replisome progression fails. These new findings reveal how a single type of helicase is recruited to the replisome to provide two independent ways of underpinning replication of protein-bound DNA, a problem that all organisms face as they replicate their genomes.

**Significance statement:** All organisms face the challenge of proteins bound to DNA acting as barriers to prevent DNA replication. We have performed fluorescence imaging experiments on living bacteria to track the positions of the replication machinery, a protein called Rep which is involved in removing these barriers, and a protein called PriC believed to be involved with reloading the replication machinery if the original replication machinery breaks down. We find that Rep is very dynamic with continual binding and movement away from the replication machinery. Association with the replication machinery depends on both binding to the replication machinery directly and on PriC. Thus Rep can circumvent barriers in two independent ways: a strategy which may be relevant to all organisms.

---

Complex multienzyme systems produce high fidelity, complete copies of genomes prior to cell division but these replisomes face frequent barriers to their continued movement along DNA, threatening genome stability. Proteins bound to the template DNA are potential barriers, with the very high stability and abundance of transcribing RNA polymerases posing a particular challenge (1, 2). Nucleoprotein barriers must be removed and replication resumed either by the original replisome or, if the blocked replisome dissociates, a replisome reloaded onto the DNA in a process known as replication restart (3). At the core of all replisomes are replicative helicases that unwind the template DNA and these replicative helicases may disrupt many, possibly most, of the potential nucleoprotein barriers encountered during genome duplication (4). However, the very high frequency of such collisions results in stochastic blockage of replisomes that requires additional mechanisms to ensure continued DNA replication (5, 6). One such mechanism uses additional helicases to promote replisome movement along protein-bound DNA in bacteria and eukaryotes (5–10). However, since loading of the hexameric replicative helicase is tightly regulated to prevent overreplication (11), recruitment of other types of helicase therefore plays an important role in promoting replisome movement through nucleoprotein complexes.

All accessory replicative helicases identified to date are members of helicase Superfamily 1, members of which translocate as monomers along single-stranded DNA either in the 5’-3’ or in the 3’-5’ direction (12). Evidence is also emerging that at least one of these accessory helicases, Rep from *E. coli*, has evolved features that optimise protein displacement from DNA (13). All accessory helicases studied so far have a polarity of translocation opposite that of the primary replicative helicase (4). Thus the *E. coli* accessory helicase Rep translocates 3’-5’ along single-stranded DNA (14) while the primary replicative helicase, DnaB, translocates 5’-3’ (15). Primary and accessory replicative helicases therefore translocate on opposing arms of the replication fork, which may allow additional motors to operate at a nucleoprotein block whilst the primary replicative helicase remains fully active at the fork (4). This arrangement may ensure that accessory helicases clear nucleoprotein barriers ahead of a paused but still active replisome that retains the primary replicative helicase, allowing resumption of replication by the same replisome without the dangers of blocked fork processing and replisome reloading (6, 8, 16–18).

Multiple monomers of Superfamily 1 helicases can function cooperatively to displace proteins from DNA (19). Having multiple accessory helicase monomers available at paused forks might therefore facilitate nucleoprotein complex removal (9). However, there is very little information concerning how accessory helicases interact physically and functionally with the replisome. The accessory helicases in *Saccharomyces cerevisiae* and *Schizosaccharomyces pombe*, Rrm3 and Pfh1 (5, 20, 21), interact with one or more subunits of the replisome (22–24). Similarly, *E. coli* Rep interacts via its C-terminus with the primary replicative helicase DnaB resulting in cooperative DNA unwinding and protein displacement by Rep and DnaB *in vitro* (6, 9, 25, 26). There is the potential for up to six Rep monomers to associate with hexameric DnaB at the *E. coli* fork, supporting a model of multiple monomer recruitment to aid protein clearance (9). However, DnaB is a protein-protein interaction hub for the entire replisome (27) and so not all of the six DnaB subunits might be accessible to Rep. Indeed, a recent live cell single-molecule imaging study failed to detect any Rep molecules present at the replisome (28). Furthermore, accessory helicases at the fork may have more than one function and more than one interaction partner. Rep may interact functionally with the replication restart protein PriC to aid replisome reloading in the event of fork stalling and replisome dissociation (3, 29, 30). Rep may unwind the nascent lagging strand at such stalled forks to expose single-stranded DNA for PriC-directed loading of DnaB back onto the fork (30). Untangling the functions of Rep in promoting fork movement along protein-bound DNA and in replication restart is difficult, though. Loss of Rep accessory helicase function results in increased fork pausing and therefore fork breakdown, leading to an increased requirement for replication restart (31). How accessory replicative helicases operate within the context of replisomes to promote genome duplication remains obscure therefore.

Here we use single-molecule microscopy of Rep in live *E. coli* cells and demonstrate that Rep colocalizes with ∼70% of replication forks. When present, there are six Rep monomers associated with each replisome, a stoichiometry that depends on the Rep-DnaB interaction, indicating maximal occupancy of the single DnaB hexamer within the replisome. Rep molecules associate only transiently with the replisome, in part due to Rep-catalysed ATP hydrolysis, indicating dynamic association with the replisome and then translocation away from the fork. PriC is also involved in co-localization of Rep with the replisome, with loss of both the Rep-DnaB interaction and PriC being required to abolish colocalization of Rep with the replication fork. There are therefore two populations of Rep associated with replisomes *in vivo*. One population might involve Rep molecules continually associating with DnaB and then translocating away to aid nucleoprotein barrier removal ahead of the fork, while the second population aids PriC-dependent reloading of DnaB in case replisome progression fails. These findings reveal for the first time the disposition of an accessory helicase within the context of a replication fork *in vivo*. They also reveal how a single type of helicase is recruited to the replisome to provide two ways of underpinning replication of protein-bound DNA, a problem that all organisms must face as they replicate their genomes.

## Results

### Rep hexamers associate with most replication forks, with monomeric Rep diffuse in the cytoplasm

We set out to test the extent of association between Rep and functional replication forks, and what mediates this interaction. To report on the replisome position we replaced the wild type *dnaQ* gene on the chromosome with a C-terminal *dnaQ-mCherry* fusion construct (see *SI Appendix)* using lambda red recombineering (32) as well as replacing either wild type copies of *rep* or *priC* genes with N-terminal monomeric GFP (mGFP) fusions (33) *mGFP-rep* and *mGFP-priC* respectively to generate two dual-label strains expressing either mGFP-Rep or mGFP-PriC, with a DnaQ-mCherry fork marker, both with wild type levels of functional activity (Fig. S1; SI Table S1). To observe the dynamic patterns of Rep and PriC localization in the cell relative to the replication fork we used single-molecule Slimfield imaging (34). This optical microscopic technique allows detection of fluorescently-labelled proteins with millisecond sampling to within 40 nm precision (35), enabling real time quantification of stoichiometry and mobility of tracked molecular complexes inside living cells, exploited previously to study functional proteins involved in DNA replication and remodelling in bacteria (36, 37), bacterial cell division (38), eukaryotic gene regulation (39), and chemokine signalling in lymph nodes (40).

**Figure 1:**
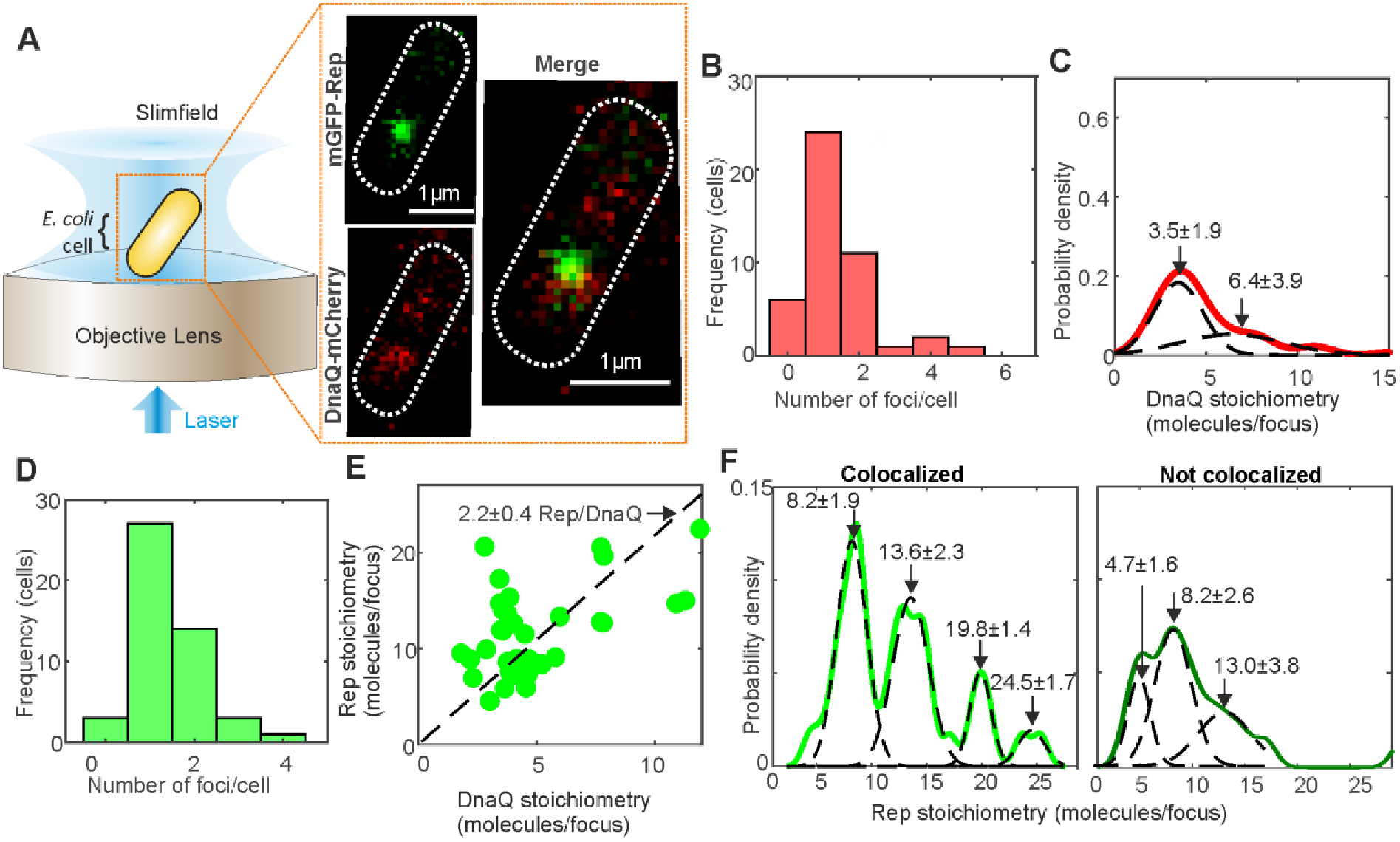
Single-molecule Slimfield of DnaQ-mCherry and mGFP-Rep. A. Slimfield schematic and example images of mGFP-Rep (green) and DnaQ-mCherry (red). B. Histogram showing DnaQ-mCherry foci detected per cell. C. Kernel density estimate of number of DnaQ-mCherry molecules per focus with two Gaussian fit, means and SEM indicated. D. Number of Rep foci per cell. E. Rep *vs*. DnaQ stoichiometry for all colocalized foci (green), linear fit constrained through origin indicated with gradient ± 95% confidence interval. F. Stoichiometry of Rep foci colocalized and not to DnaQ, multiple Gaussian fits shown with mean ± SD indicated. N=45 cells.

We grew cells to mid-logarithmic phase then immobilized cells onto agarose pads suffused with growth medium for imaging. Slimfield indicated mostly one or two replication forks per cell (Fig. 1A), manifest as distinct fluorescent foci of diffraction-limited width ∼300 nm, as expected for cells undergoing mainly one round of chromosomal duplication per cell cycle as we have in our growth conditions (36). Using step-wise photobleaching analysis of the mCherry tag we could accurately quantify the stoichiometry of these foci (Fig. S2) indicating peaks centered on three or six DnaQ molecules per focus (Fig. 1C) to be compared with previous observations from live cell fluorescence microscopy (36, 41) indicating three DNA polymerases per replisome (42, 43), or six per focus when two replication forks are sufficiently close so that they cannot be resolved optically. Replacing the fluorophore with mGFP (Fig. S3) yielded similar stoichiometries but with more foci detected per cell consistent with its smaller point spread function width and higher emission signal relative to mCherry (37).

In the same DnaQ-mCherry containing cells we observed mostly one or two mGFP-Rep foci per cell (Fig. 1D). By computing the numerical overlap integral between foci in the red and green detection channels (44) we could robustly determine the extent of colocalization between Rep and DnaQ to within 40 nm localization precision. These analyses indicated that 70±7% (±SE, N=70 foci) of Rep foci were colocalized with DnaQ, with both the colocalized and non colocalized populations having similar ranges of stoichiometry equivalent to approximately 6–30 Rep molecules per focus, however, only colocalized Rep displayed distinct periodicity in stoichiometry; a linear fit of Rep to DnaQ stoichiometry showed approximately two Rep molecules associated per DnaQ molecule (Fig. 1E, Fig. 1F). Since each replisome contains an average of three DnaQ molecules (36, 42) our data indicate there are an average of six Rep molecules present at each replisome, consistent with our measurement of the periodicity of Rep stoichiometry colocalized with DnaQ (Fig. 2E). An average of six Rep molecules associated with each replisome indicates full occupancy of Rep binding sites on the DnaB hexamer within each replisome (9).

**Figure 2:**
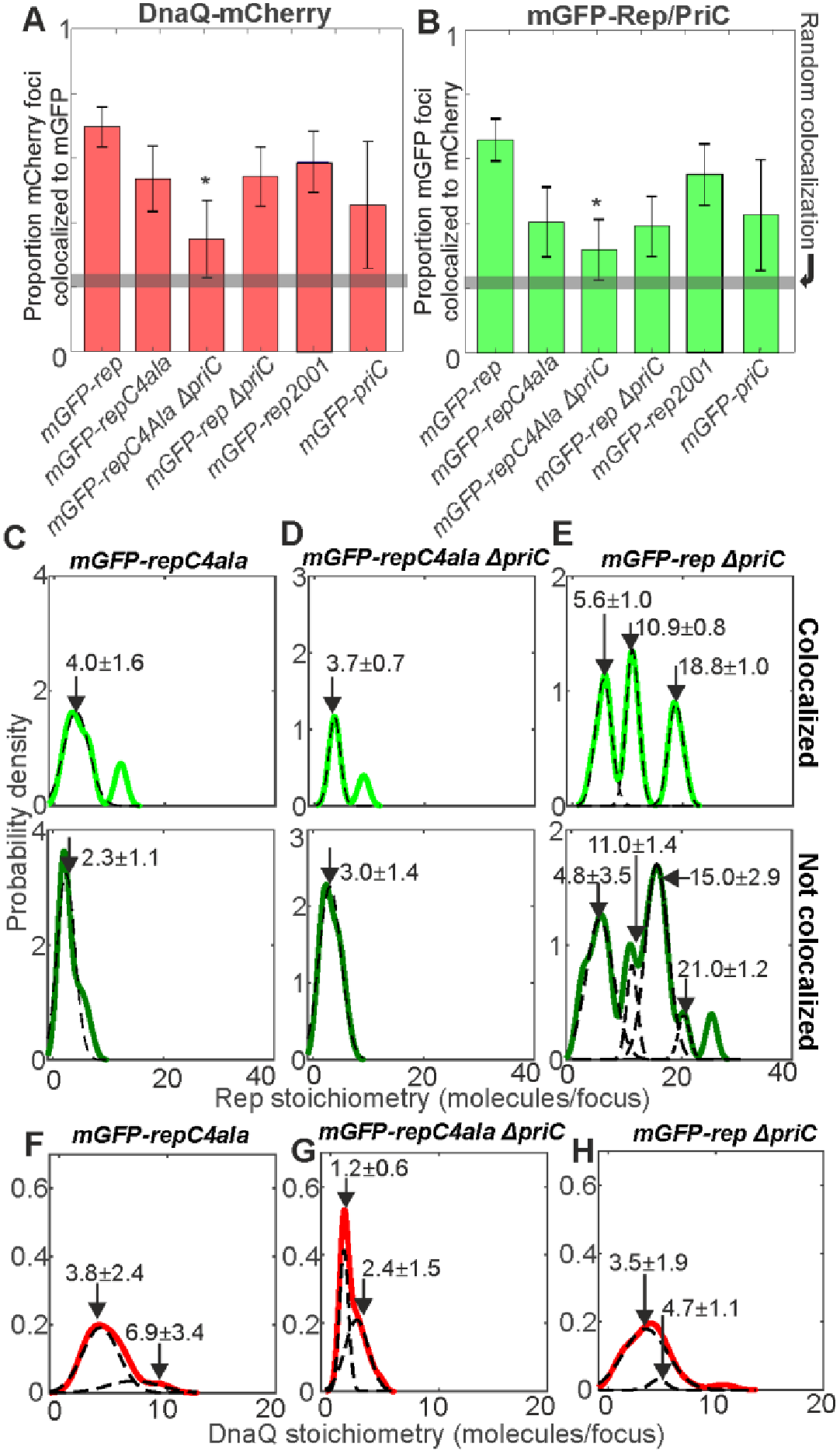
Rep/DnaQ colocalization analysis. Proportions of A. DnaQ-mCherry foci colocalized with mGFP-Rep or mGFP-PriC, B. mGFP-Rep or mGFP-PriC foci colocalized with DnaQ-mCherry. All strains carry *dnaQ-mCherry* allele with relevant genotypes indicated, gray horizontal bar indicates random colocalization level based on our simulations. C-E. Kernel density estimates of number of mGFP-Rep molecules in foci colocalized with DnaQ-mCherry (light green) and foci not colocalized with DnaQ-mCherry (dark green) in wild type and mutant backgrounds. Multiple Gaussian fits (dotted lines) and mean values ± SD indicated. F-H. Kernel density estimates of the number of DnaQ-mCherry in each focus in wild type and mutant backgrounds, multiple Gaussian fits (dotted lines) and mean values ± SD indicated. N=30 cells.

As well as distinct foci, we also detected a diffuse pool of Rep fluorescence throughout the cell, similar to previous studies of *E. coli* replisome proteins (36). Using numerical integration of cellular pixel intensities (45) we quantified the pool copy number to be several hundred Rep molecules per cell (Fig. S7) comparable to that estimated previously using quantitative western blots on cell lysates (46).-We can estimate the stoichiometry of Rep foci in the pool using nearest neighbor analysis (39), since by definition pool foci must be separated by less than the optical resolution limit of our microscope which is ∼230 nm, indicating monomeric Rep in the pool (see *SI Appendix)*.

### Rep-fork association is mediated by DnaB

The Rep-DnaB interaction resides within the C-terminal 33 amino acids of Rep and consequently the *repΔC33* allele displays a partial loss of *rep* function (9, 46). To test whether the patterns of colocalization between Rep and the replisome we observed were due to the Rep-DnaB interaction we constructed an *mGFP-repΔC33* fusion. However, the fusion had a negative impact on *repΔC33* function (Fig. S5C compare iii with iv). We therefore searched for mutations within the C-terminal 33 codons that would recapitulate the *repΔC33* phenotype but would otherwise retain function when fused to *mGFP*. We found that mutating the final four codons of *rep* encoding KRGK to encode alanine resulted in an allele displaying a partial loss of function similar to *repΔC33* but which could be fused to *mGFP* without a complete loss of function (Fig. S5; Table S1).

When *mGFP-repC4ala* was introduced into the *dnaQ-mCherry* strain the fraction of colocalized Rep-DnaQ foci dropped significantly (Fig. 2A and B) but similar numbers of foci were detected (Fig. S6). The stoichiometry of RepC4Ala foci dropped to 2–4 molecules per focus independent of their position relative to the fork, however, we observed that fork-colocalized RepC4Ala foci lost the pattern of periodicity in the stoichiometry distribution which we observed with mGFP-Rep (compare Fig. 2C with 1E), suggesting a key role in the Rep C terminus in determining its hexameric structure when associated with DnaB. However, levels of colocalization seen with RepC4Ala were still above those expected for purely random optical overlap of Rep and DnaQ foci (Fig. 2A and B). This non-random association indicates either that RepC4Ala can still interact with DnaB to some extent or that Rep can associate with the replisome independent of the Rep-DnaB interaction. The significant decrease in the number of RepC4Ala molecules within foci that are not colocalized with DnaQ as compared with wild type Rep (compare Fig. 2C with Fig. 1E) also indicate that the Rep C-terminus plays a role in the formation of Rep oligomers in the absence of any direct association with the replication fork.

### Association of Rep and replication forks is modulated by PriC

Biochemical and genetic evidence indicates that Rep also participates in PriC-dependent fork reloading (29, 30). However, evidence of a physical association between PriC and Rep is lacking, prompting us to employ functional imaging of PriC in live cells. Using an *mGFP-priC* fusion that retained wild type function (Figure S1B), we found that ∼40% of DnaQ foci contained PriC (Fig. 2A and S4). Thus a significant minority of replisomes contain PriC.

The impact of PriC on the colocalization of DnaQ and Rep was probed by deleting *priC*. A *ΔpriC* mutation reduced the proportion of Rep-DnaQ colocalized foci (Fig. 2B). The probability of Rep association with the replisome is therefore determined in part by PriC. In contrast, the range of stoichiometries of Rep molecules in foci colocalized with DnaQ was relatively unaffected when comparing *priC^+^* and *ΔpriC* strains, and the hexameric periodicity in stoichiometry remained (compare Fig. 2E and 1E), which contrasts with the marked impact of the *repC4ala* mutation. These data indicate that the pronounced periodicity in the patterns of association of Rep with the replisome is dependent on the Rep C-terminus rather than PriC.

Combining both *repC4ala* and *ΔpriC* mutations reduced the incidence of RepC4Ala colocalization with DnaQ to levels consistent with random association with the replisome (Fig. 2A and B). Thus both the Rep C-terminus and PriC contribute to association of Rep with the replisome. However, the stoichiometry of RepC4Ala foci associated with DnaQ in the *repC4ala ΔpriC* double mutant strain was similar to the single *repC4ala* mutant (Fig. 2, compare C and D). The significant periodicity in patterns of association of Rep with the replisome is determined therefore by the Rep C-terminus rather than PriC. Replisome composition was also affected in the *repC4ala ΔpriC* double mutant since the number of DnaQ molecules was reduced from 3–6 to 1–2 molecules per focus (compare Fig. 2D with 1E).

Deleting *priC* also altered the pattern of Rep stoichiometry in foci not colocalized with the replisome (compare Fig. 1G with Fig. 2E). However, there were still significant numbers of Rep molecules in foci far from the replisome in *ΔpriC* cells which was in marked contrast to the major reduction in numbers of RepC4Ala molecules in foci far from the replisome in *priC^+^* cells (compare Figure 2C and E). These data indicate that the Rep C-terminus is the primary determinant of Rep oligomer formation far from the replisome, as with focus formation at the replisome.

### Rep-fork interactions transient, dynamic and ATP dependent

The generally accepted model of Rep accessory helicase function is that Rep associated with the replisome translocates along the single-stranded leading strand template and unwinds the parental dsDNA whilst simultaneously promoting dissociation of any proteins bound to this dsDNA (9) (see also Fig. 4). Rep might therefore translocate in an ATP-dependent manner away from the replisome in addition to any spontaneous dissociation. We probed therefore the ATP dependence of Rep-DnaQ dissociation, and its dynamics.

Rep foci appeared highly dynamic (Supplementary Movie 1 and Movie 2). We analysed their mobility on the millisecond timescale, correlated to their state of localization with the fork, by calculating the microscopic diffusion coefficient *D* of each tracked focus and fitting a model consisting of the sum of multiple gamma functions model (40). A three parameter models fitted the data best (Fig. 3A and B and Fig S8), comprising *D* = 0.09 μm^2^/s, consistent with immobile foci based on our tracking localization precision of 40 nm, in addition to a slow (D = 0.4 μm^2^/s) and a fast (D =1.3 μm^2^/s) diffusion mode.

**Figure 3.**
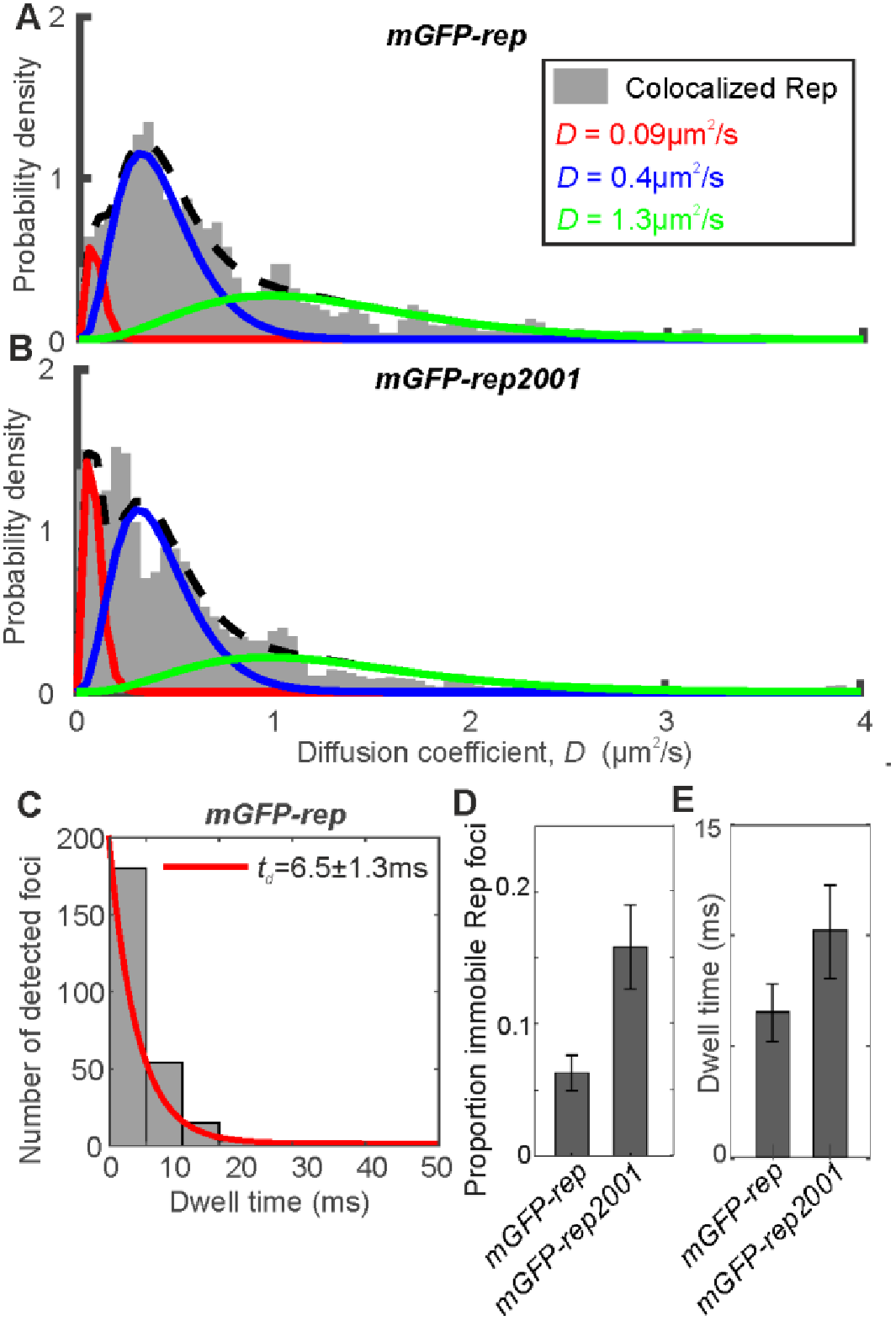
Rep mobility analysis. A and B. Binned kernel density estimates (gray) of *mGFP-rep* and *mGFP-rep2001* diffusion coefficient distributions with three Gamma curve diffusion coefficient fits, minimal reduced χ^2^=0.0067, proportion in each model indicated. C. mGFP-Rep foci dwell time with mCherry-DnaQ foci distribution with an exponential fit (red). D. Proportion of colocalized Rep foci that are immobile, as determined from the three Gamma curve fits. E. Histogram for the distribution of mean dwell time derived from fits for *mGFP-rep* and *mGFP-rep2001*. Error bars are 95% confidence intervals.

To probe the dependence on ATP hydrolysis we labeled a mutant RepK28R with mGFP, whose mutation lies in the Walker A domain that is essential for ATP hydrolysis and hence translocation along DNA (46, 47). The mGFP-RepK28R fusion retained the ability to associate with DnaQ, as evidenced by a similar proportion of colocalized mGFP-RepK28R and DnaQ foci as compared with mGFP-Rep (Fig. 2B). Also, the distributions of RepK28R stoichiometry (Fig. S6) and total cell copy number (Fig. S7) were similar to wild type. However, mGFP-RepK28R also showed a significant increase in the proportion of immobile colocalized foci from 6±1% in the wild type to 15±3% (compare Fig. 3B with 3A; Fig. 4D). This increase contrasted with Rep foci not colocalized with the fork, which failed to show a significant difference between wild type and RepK28R (Fig. S8C).

**Figure 4.**
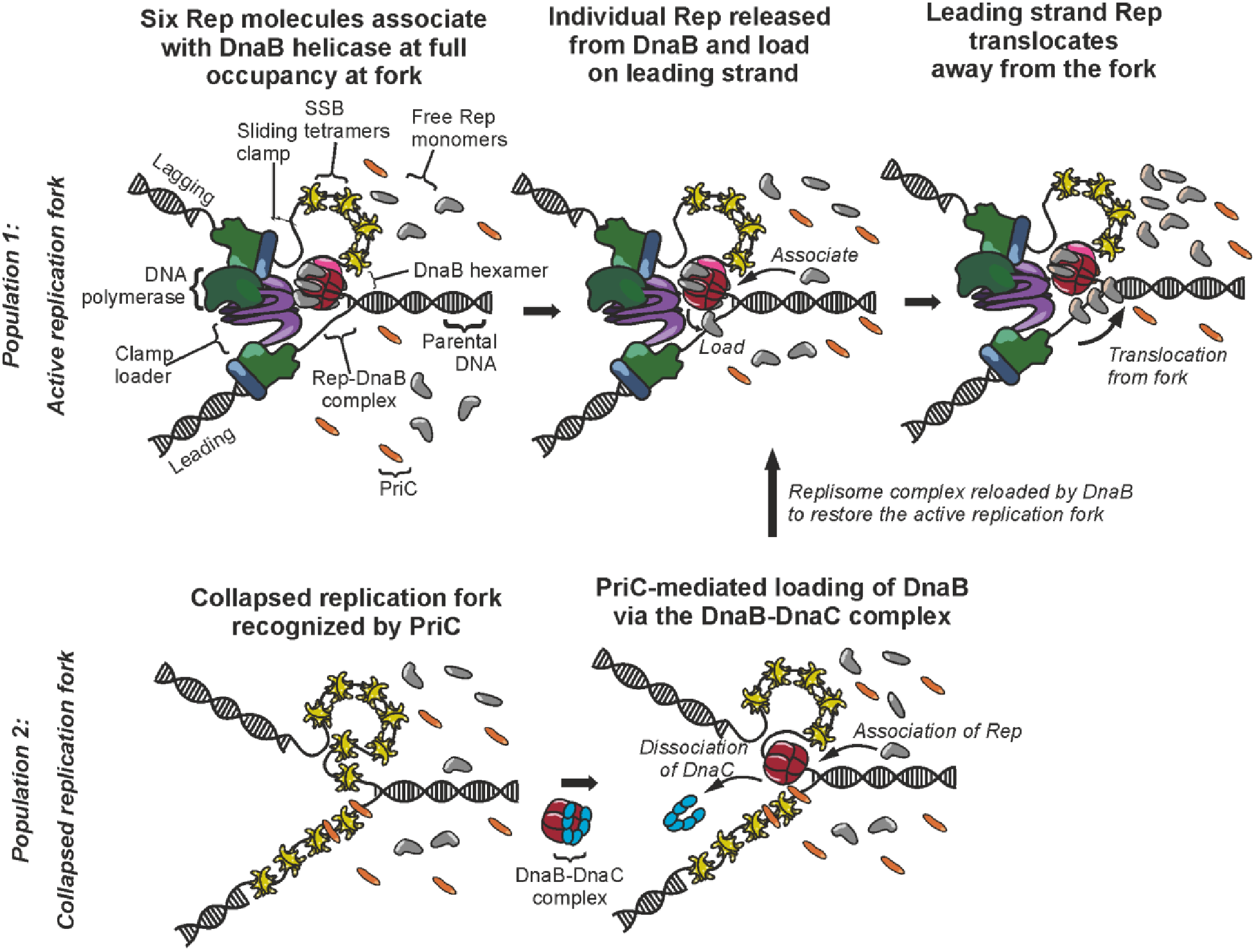
Model representing the two populations of Rep interactions. Population1: Interaction of Rep with replicative helicase DnaB. Monomers of Rep (gray) associate with individual monomer subunits within the DnaB (red) hexamer to full occupancy of the hexamer. Rep monomers are continuously released from DnaB and load onto the leading strand. Released Rep monomers then translocate from the fork coupled to the hydrolysis of ATP. Additional Rep monomers from the cytoplasm are continuously recruited onto the DnaB hexamer as vacant binding sites become available. Rep can associate with PriC to stimulate replisome reloading. Population 2: A collapsed replication fork is recognized by PriC. DnaB is then loaded via the DnaB-DnaC complex. *Legend: DNA polymerase complex - green; sliding clamp - blue; clamp loader complex - purple; DnaB - red; DnaG - pink; single-strand binding protein (SSB) - yellow; Rep - gray; PriC-orange; DnaC-cyan*.

We estimated the dwell time of Rep foci at the replication fork from the number of consecutive image frames associated with each colocalized track. The distribution of dwell times decreased exponentially with a characteristic time constant of 6.5±1.3 ms at the fork for wild type Rep, increasing to 10.2±2.1 ms with RepK28R (Fig. 3C and 3E, and S8). Dwell time fits to the *repC4ala* mutation and *priC* deletion based on a single exponential model were poor suggesting that there are likely to be a range of factors influencing dwell time, for example the kinetics of binding to and unbinding from single-stranded DNA and the frequency with which single-stranded DNA regions become available and accessible, which we propose to investigate in future studies. We conclude that when Rep is able to hydrolyze ATP, a smaller proportion of Rep molecules are immobile at the replisome and these immobile molecules also spend significantly less time at the fork. These data imply that dissociation of Rep from the replisome is driven in part by ATP-dependent translocation of Rep along DNA.

## Discussion

Here we show that the majority of replisomes contain the accessory replicative helicase Rep, that there are approximately six Rep molecules per replisome and that this distribution is dependent upon the Rep C-terminus (Fig. 1 and 2). These data are consistent with Rep association being driven primarily by the Rep-DnaB interaction (9) and indicate high occupancy of the six Rep binding sites within the DnaB hexamer at the replisome. Our data also demonstrate rapid turnover of Rep at the replisome and the importance of Rep-catalysed ATP hydrolysis for this rapid turnover (Fig. 3). These findings suggest a model in which the majority of replisomes have near-full occupancy of Rep binding sites and that these Rep molecules bind continually to single-stranded DNA at the fork to translocate ahead of the advancing replisome to help displace proteins from the template. We also find that association of Rep with the replisome is dependent in part on PriC (29, 30) (Fig. 2B), consistent with the functional interaction between Rep and PriC in replication restart. Whether this PriC-dependent association of Rep with the replisome is due to a direct Rep-PriC interaction or due to an indirect effect of PriC is unknown. These data do indicate, though, that there may be a complex interplay between DnaB and PriC in terms of Rep function within the replisome.

Our data also demonstrate that a minority of Rep foci form away from any replisomes (Fig. 2B and 1F) with the number of Rep molecules within these foci dependent primarily on the Rep-DnaB interaction (compare Fig. 1F with Fig. 2C). DnaB hexamers can be loaded onto single-stranded DNA only with the aid of the helicase loader DnaC (48–51) implying that at least some of the DnaB not associated with replisomes is bound by DnaC in a DnaB_6_:DnaC_6_ complex (52). Our data indicate that at least some of this DnaB not within replisomes is associated with Rep, consistent with earlier observations for live cell fluorescence microscopy that mobile DnaB foci can be detected diffusing away from replication forks in addition to an immobile replisome-anchoring population (53). The binding of Rep and DnaC to DnaB appears to be mutually exclusive (9) implying that Rep and DnaC are in competition for binding of the pool of DnaB away from replisomes.

Our finding of multiple Rep molecules colocalized with the replisome compares to a previous recent live cell imaging study of fluorescently-labeled Rep, and other repair and replisome proteins (28). Here, although the authors did not have an independent fork marker for visualizing simultaneous Rep and fork colocalization, they observed Rep foci in locations consistent with fork localization. They reported populations of Rep foci which were relatively stable in appearing in at least four consecutive image frames, but also a significant number of foci that lasted for fewer than four consecutive frames. The total proportion of Rep foci in locations consistent with association of the replication was ∼70% (comprising 32% stable and 38% unstable foci), similar to the proportion which we report here from our more direct approach using an independent fork marker. Our observations are consistent with these previous findings in light of the very rapid dynamics of Rep we measure (average dwell time of ~6 ms at the fork) which is significantly faster than the earlier study could sample with a frame integration time of 40 ms strobed every 200 ms. Coupled to this Rep foci detection in this earlier study was also limited to a reported sensitivity of at best 3–4 fluorescent protein molecules per immobile focus, but likely to be substantially worse for Rep due to blurring of the fluorescent protein optical image in light of the rapid dynamics at the fork, which taken together explains the apparent appearance of lower stability foci reported in the earlier study.

What are the implications of our data for the functioning of Rep as an accessory replicative helicase? Our data are consistent with Rep molecules bound to the DnaB hexamer associating continually with single-stranded DNA at the fork and translocating along this ssDNA in an ATP-dependent manner away from the replication fork. The 3’-5’ polarity of Rep translocation along ssDNA and the occlusion of the lagging strand template by the DnaB hexamer makes it likely that any Rep translocation will be along the leading strand template, consistent with Rep movement along this strand ahead of the fork to displace proteins out of the path of the advancing replisome (4, 9). Such a model implies that at the majority of replisomes there is a continual firing of Rep molecules ahead of the replisome, analogous to bullets in a revolver. Having multiple Rep molecules translocating ahead of the fork might be needed for effective unwinding of double-stranded DNA and hence protein displacement ahead of the fork, given the inability of Rep monomers to unwind DNA *in vitro* in the absence of partner proteins (54).

How does this model of accessory helicase activity interface with Rep acting as an accessory factor in PriC-catalysed reloading of DnaB onto the lagging strand template during replication restart? The significant periodicity of numbers of Rep associated with the replisome depends on the Rep C-terminus rather than PriC (compare Fig. 1F with Fig. 2C and E), consistent with Rep association with the replisome being dominated by the Rep-DnaB interaction. However, the presence of PriC at 40% of forks leads to additional Rep molecules being associated with the replisome (Fig. 2B). Colocalization of Rep with the replisome depends therefore upon both the Rep C-terminus and on PriC (Fig. 2B). There are therefore two pools of Rep at the replisome, one pool dependent upon the Rep-DnaB interaction and another pool dependent on PriC (Fig. 4). PriC interacts with single-stranded DNA and with SSB (55, 56) providing means by which PriC could interact with the replisome and hence recruit Rep. Evidence for a direct Rep-PriC interaction is currently lacking but it is also possible that PriC recruits Rep to the replisome indirectly. Both PriC and Rep also interact with DnaB (9, 25, 57), and so association of PriC with DnaB might result in allosteric effects on DnaB that affect the known Rep-DnaB interaction (31). However, PriC is responsible for some colocalization of Rep with the replisome even in the Rep C-terminal mutant (Fig. 2B) indicating that PriC-dependent recruitment of Rep is likely to be independent of any Rep-DnaB interaction.

Regardless of whether our observed PriC-dependent association of Rep with the replisome is a direct or indirect effect, our data lend support to a functional Rep-PriC interaction inside cells (29, 30). Different dispositions of DnaB and PriC with respect to DNA within the replication fork might facilitate two different functions for the two different pools of Rep at the replisome. The DnaB-dependent pool of Rep very likely promotes replisome progression along protein-bound DNA via translocation of Rep along the leading strand template ahead of the fork (9). The second pool, associated directly or indirectly with PriC, might aid PriC-directed reloading of DnaB back onto the fork via Rep-catalyzed unwinding of the lagging strand duplex at the fork to generate single-stranded DNA for DnaB binding (30). Recruitment of Rep by two different factors at the replisome might therefore provide two ways in which Rep facilitates duplication of protein-bound DNA. However, the interplay between Rep and PriC is difficult to resolve. While PriC provides a pathway of replication restart, the accessory helicase function of Rep reduces the need for replication restart, complicating interpretation of this interplay. Our data do indicate, though, the importance of Rep and PriC for maintaining the architecture of the replisome. In both *repC4ala priC^+^* and *rep^+^ ΔpriC* cells the number of DnaQ molecules per focus is on average three, as found in wild type cells (36) (compare Fig. 1C with Fig. 2F and H). However, *repC4ala ΔpriC* cells have only 1–2 DnaQ molecules per focus (Fig. 2G). This reduction in DnaQ molecules at the replisome is unlikely to be due to allosteric effects upon the structure of replisomes lacking Rep and PriC since replisomes that lack both Rep and PriC *in vitro* retain three DnaQ molecules (42). Alternatively this altered replisome architecture might be due to increased pausing and blockage of the replisome at nucleoprotein barriers in the absence of an accessory replicative helicase coupled with defective replisome reloading without PriC. Regardless of the reasons for this altered replisome structure, our data indicate that both Rep and PriC are important constituents of the replisome.

## Materials and Methods

### Cell strains

*E. coli* clones comprising fluorescently tagged alleles of the *dnaQ, rep*, and *priC* genes (SI Table S2) were introduced into the respective native loci by lambda red recombineering (32), full details of cell doubling (SI Table S1), plasmids (SI Table S3), primers (SI Table S4) and methods in SI Appendix. Cells were routinely grown overnight in LB at 37°C from freshly streaked LB plates. The LB grown cultures were then subcultured to mid-log phase at 30°C in 1X 56 salts minimal medium with 0.2% glucose as the carbon source. The cells were then spotted onto slides overlaid with 1% agarose containing 1X 56 salts and 0.2% glucose.

### Microscopy and image analysis

A Slimfield microscope was used (39) for single-molecule imaging, an Olympus BX63 microscope measured epifluorescence. Foci tracking used MATLAB (Mathworks) software which determined *D* and stoichiometry using foci brightness {Miller, 2015 #33587} and Chung-Kennedy (58) filtered mYPet step-wise photobleaching, and nearest-neighbor modeling (44). Full details SI Appendix.

### Data availability

Data included in full in main text and supplementary files. Raw data available from authors.

### Software access

Code written in MATLAB available at https://sourceforge.net/projects/york-biophysics/.

## Acknowledgments

Supported by BBSRC (grant BB/N006453/1) to P.M. and M.L. Part-funded by the Wellcome Trust [ref: 204829] through Centre for Future Health at the University of York to A.J.M.W.

## Supplementary Appendix

**SI Table S1.** Doubling time of labeled strains. Three replicates performed for each strain, OD measurements binned into 10 min interval time points.

| Strain | Mean doubling time (min) | SD (min) | SEM (min) |
| --- | --- | --- | --- |
| WT | 80 | 10 | 5.8 |
| *dnaQ-mGFP-<kan>* | 90 | 10 | 5.8 |
| *dnaQ-mCherry-<kan>* | 93.3 | 5.8 | 3.3 |
| *mCherry-rep-<kan>* | 96.7 | 5.8 | 3.3 |
| *mGFP-rep-<kan>* | 93.3 | 5.8 | 3.3 |
| *mGFP-repC4ala-<kan>* | 113.3 | 5.8 | 3.3 |
| *mGFP-rep2001-<kan>* | 113.3 | 11.5 | 6.7 |
| *dnaQ-mCherry-<> mGFP-rep-<kan>* | 96.7 | 11.5 | 6.7 |
| *dnaQ-mGFP-<kan> mCherry-rep-<>* | 96.7 | 11.5 | 6.7 |
| *mGFP-priC-<kan>* | 103.3 | 5.8 | 3.3 |
| *dnaQ-mCherry mGFP-priC-<kan>* | 123.3 | 5.8 | 3.3 |

**SI Table S2:** Strains used in this study.

| Strain | Relevant genotype | Source or derivation |
| --- | --- | --- |
| BW25113 derivatives: |  |  |
| BW25113 | *rrnB* Δ*lacZ*4787 *hsdR*514 $\Delta$(*araBAD*)567<br>$\Delta$(*rhaBAD*)568 *rph*-1 | (1) |
| JW0456 | Δ*priC*752::*kan*> | (1) |
| JW5604 | Δ*rep*729::*Kan*> | (1) |
| TB28 derivatives: |  |  |
| TB28 | MG1655 Δ*lacI*ZYA | (2, 3) |
| N7150 | $\Delta$ *lacI* ZYA:: *<>* *rpoB* *35 *uvrD*:: *dhfr*<br>$\Delta$ *rep*:: *cat* / pAM407 (*uvrD* <sup>+</sup> *lacZ* <sup>+</sup>, a pRC7<br>derivative) | (4) |
| HB139 | Δ*priC*::*<kan>* | TB28 x P1(JW0456) to Km<sup>R</sup> |
| MKG3 | Δ*priC*::*<>* | HB139 to Km<sup>S</sup> with pCP20 |
| AM2017 | Δ*lacI*ZYA Δ*priB*::*dhfr* | (5, 6) |
| JGB129 | Δ*priC* / pKD46 | MKG3 transformed with pKD46 |
| JGB161 | Δ*priB*::*dhfr* / pAM421 (*priC*<sup>+</sup> *lacZ*<sup>+</sup>, a pRC7<br>derivative) | AM2017 transformed with pAM421 |
| JGB167 | *ΔpriB ΔpriC* / pAM421 (*priC<sup>+</sup> lacZ<sup>+</sup>*, a pRC7 derivative) | JGB161 x P1(JW0456) to Km<sup>R</sup> |
| JGB255 | *Δrep::apra* | *apra* integrated into AS97 |
| JGB257 | *Δrep::apra* / pKD46 | JGB255 transformed with pKD46 |
| JGB264 | *mCherry-rep <Kan>* | *mCherry-rep <Kan>* recombineered into JGB257 |
| JGB265 | *mCherry-rep-<kan>* | TB28 x P1(JGB263) to Km<sup>R</sup> |
| JGB266 | *mCherry-Rep <Kan>* | TB28 x P1(JGB264) to Km<sup>R</sup> |
| JGB286 | *mCherry-Rep <>* | JGB266 to KmS with pCP20 |
| JGB294 | *dnaQ-mGFP-<kan> mCherry-rep-<>* | JGB286 x P1(AS217) to Km<sup>R</sup> |
| KM269 | *MG1655 ΔlacIZYA::<> rpoB\*35*<br>*uvrD::dhfr Δrep729::<Kan>* / pAM407<br>(*uvrD<sup>+</sup> lacZ<sup>+</sup>*, a pRC7 derivative) | N7150 x P1(JW5604) to Km<sup>R</sup> |
| AS97 | TB28/pKD46 | TB28 transformed with pKD46 |
| AS217 | *dnaQ-mGFP-<kan>* | *mGFP-<kan>* with homology to dnaQ recombineered into AS97 |
| AS446 | *dnaQ-mCherry-<kan>* | *mCherry-<kan>* with homology to dnaQ recombineered into AS97 |
| AS448 | *dnaQ-mCherry-<>* | AS446 to km<sup>S</sup> with pCP20 |
| AS461 | *ΔlacIZYA mGFP-priC-<kan>* | *mGFP-priC-<kan>* recombineered into JGB129 |
| AS476 | JGB161 *mGFP-priC-<kan>* | JGB161 x P1(AS461) to Km<sup>R</sup> |
| AS488 | *dnaQ-mCherry-<> mGFP-priC-<kan>* | AS448 x P1(AS461) to Km<sup>R</sup> |
| AS505 | JGB257 *mGFP-rep-<kan>* | *mGFP-rep-<kan>* recombineered into JGB257 |
| AS525 | *mGFP-rep-<kan>* | TB28 X P1(AS505) to Km<sup>R</sup> |
| AS545 | *dnaQ-mCherry-<> mGFP-rep-<kan>* | AS448 X P1(AS505) to km<sup>R</sup> |
| AS554 | *dnaQ-mCherry-<> mGFP-rep-<>*<br>*ΔuvrD::dhfr* | AS545 X P1(KM269) to Tp<sup>R</sup> |
| AS609 | *mGFP-repC4ala-<kan>* | *mGFP-repC4ala-<kan>* recombineered into JGB257 |
| AS615 | *dnaQ-mCherry-<> mGFP-repC4ala-<kan>* | AS448 X P1(AS609) to Km<sup>R</sup> |
| AS662 | *dnaQ-mCherry-<> mGFP-rep-<>*<br>*ΔpriC::<kan>* | AS630 X P1(JW0456) to Km<sup>R</sup> |
| AS666 | *dnaQ-mCherry-<> mGFP-repC4ala-<>* | AS615 to km<sup>S</sup> with pCP20 |
| AS672 | *dnaQ-mCherry-<> mGFP-repC4ala-<>*<br>*ΔpriC::<kan>* | AS666 X P1(JW0456) to Km<sup>R</sup> |
| AS674 | *mGFP-rep2001-<kan>* | *mGFP-rep2001-<kan>* recombineered<br>into JGB257 |

**SI Table S3.** Plasmids used in this study.

| Plasmid | Description | Antibiotic | Reference |
| --- | --- | --- | --- |
| pCP20 | Yeast Flp recominase expression plasmid | Amp | (7) |
| pDHL580 | pUC19 *linker-mGFPmut30<kan>* | Amp, kan | (8) |
| pKD46 | $\lambda$ Red recombinase expression plasmid | Amp | (7) |
| pAS65 | pUC18 *mGFP-priC-<kan>* | Amp, kan | This study |
| pAS79 | pUC18 *mGFP-rep-<kan>* | Amp, kan | This study |
| pAS124 | pUC18 *mGFP-repC4ala-<kan>* | Amp, kan | This study |
| pAS127 | pUC18 *mGFP-rep2001-<kan>* | Amp, kan | This study |
| pJGB374 | pUC18 *linker-mCherry-<kan>* | Amp, kan | This study |
| pJGB380 | pUC18 *mCherry-rep-<kan>* | Amp, kan | This study |

**SI Table S4.**
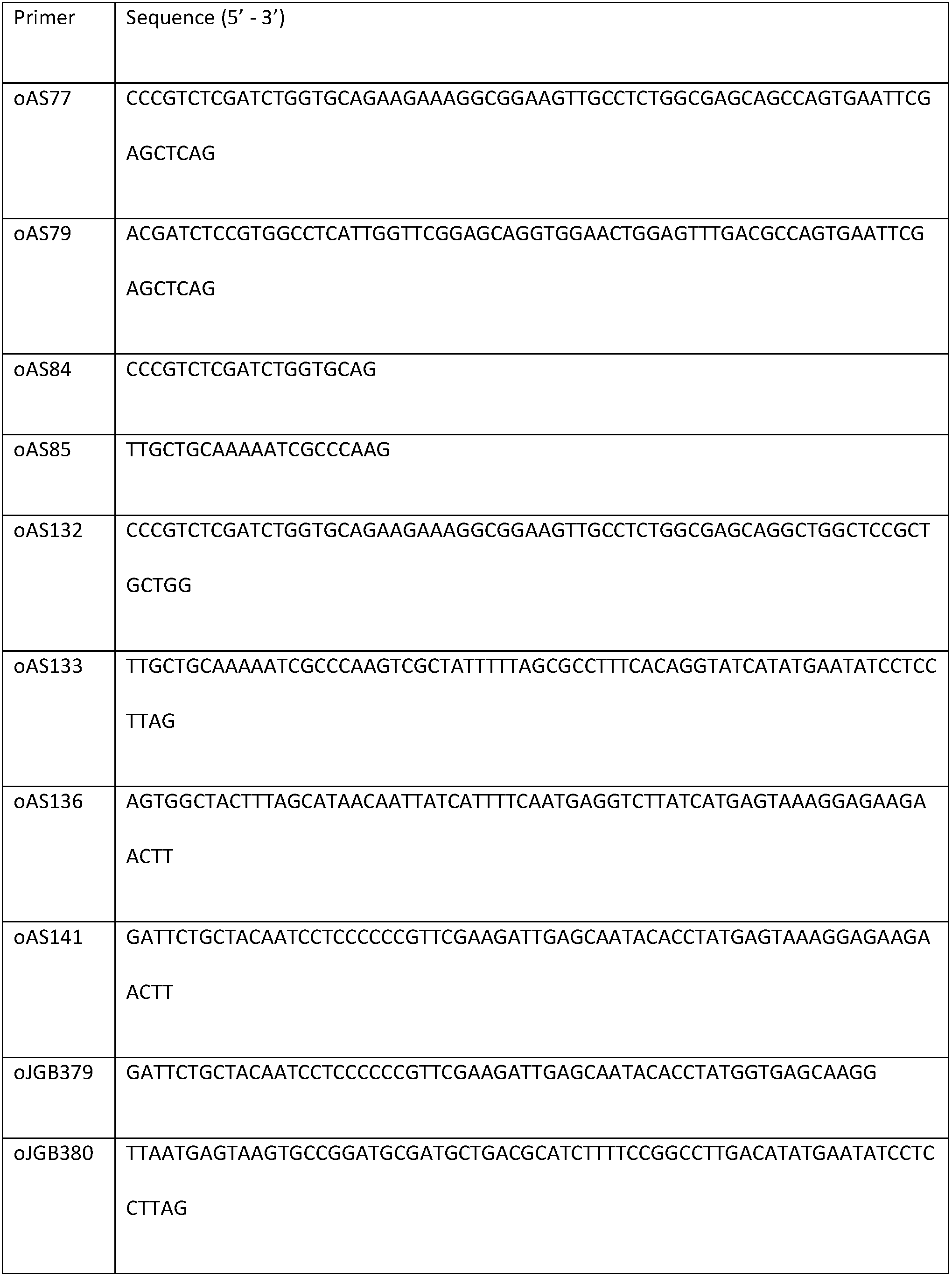
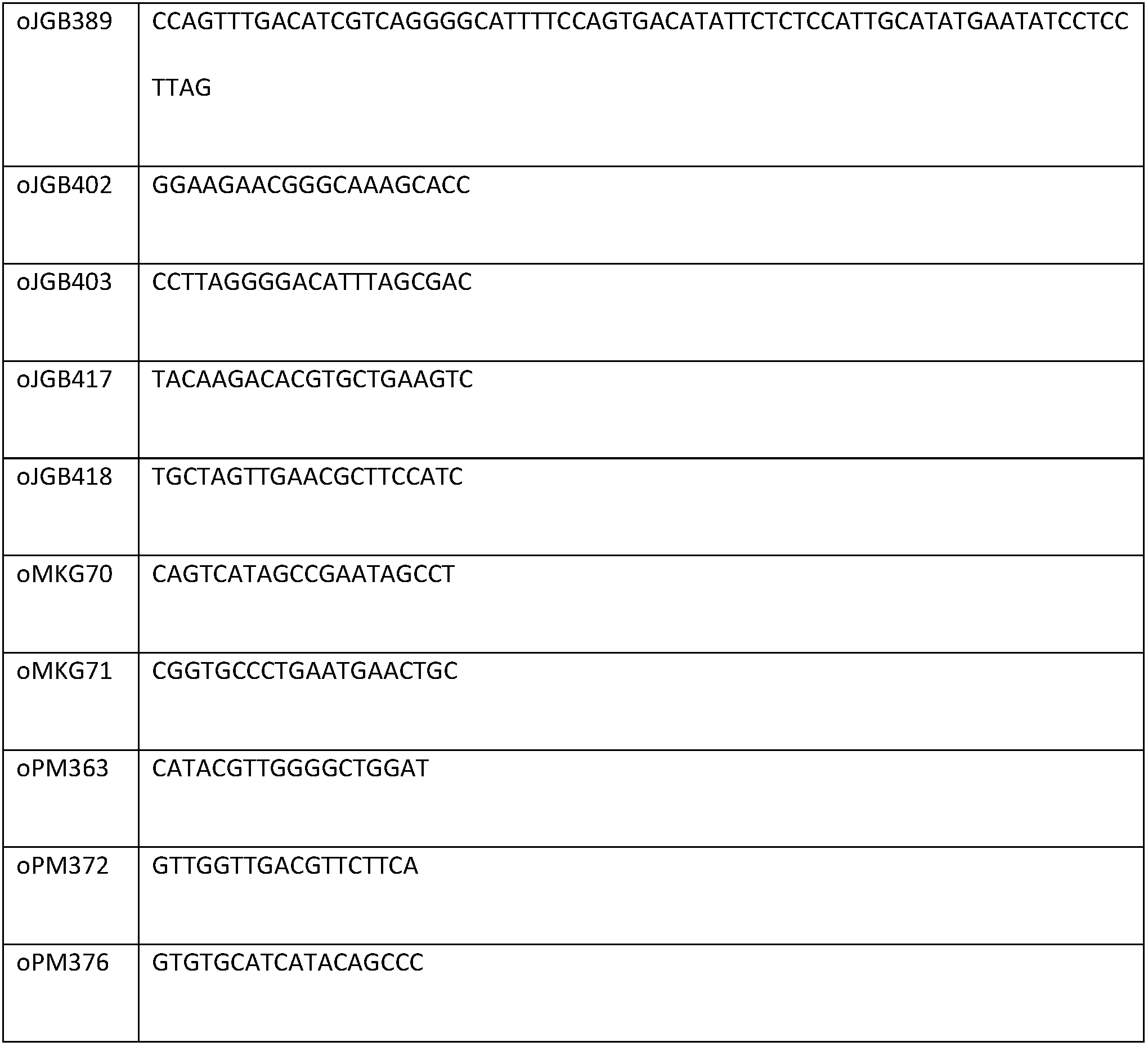
Primers used in this study.

**Figure S1:**
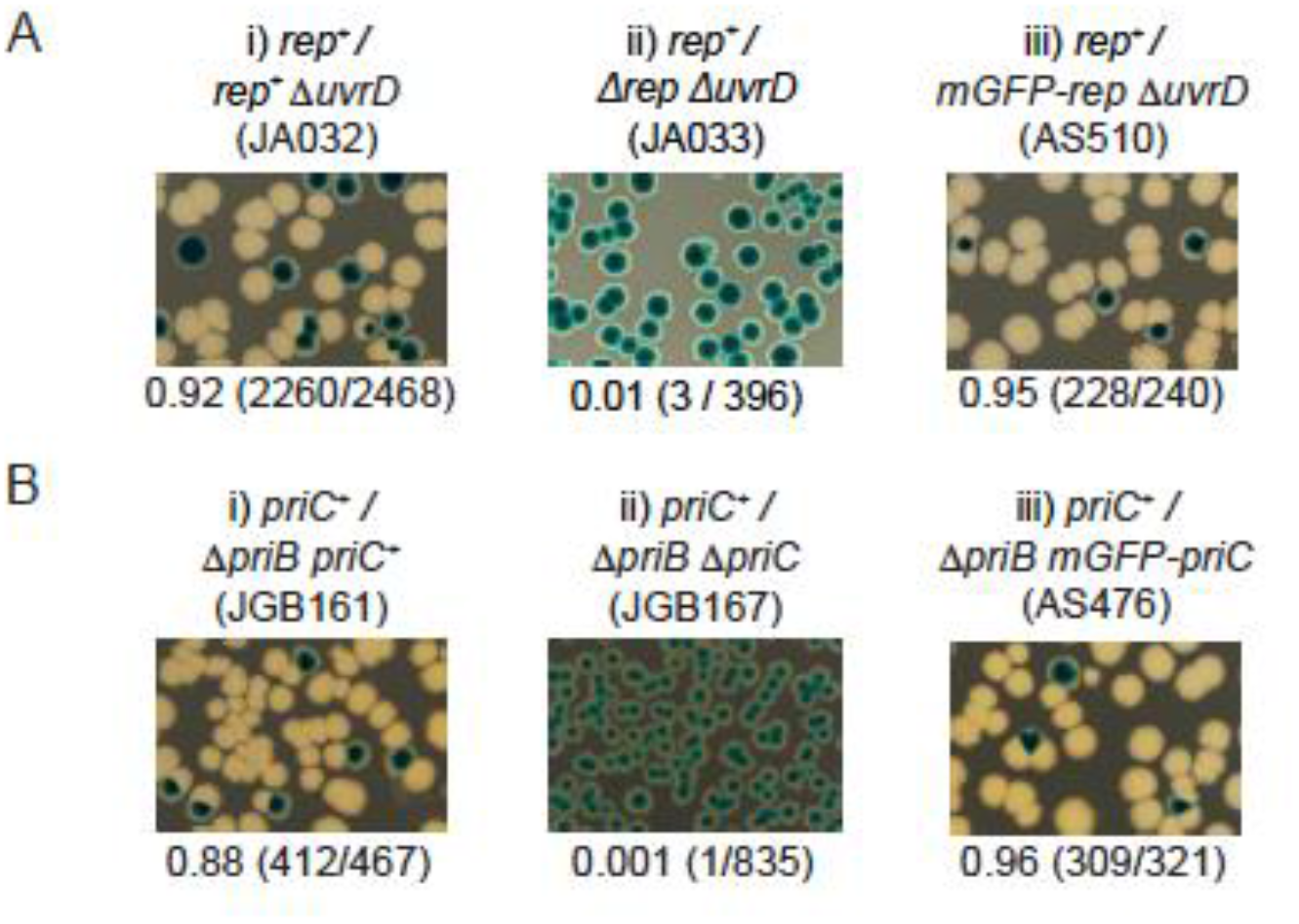
Testing of mGFP-Rep and mGFP-PriC fusions for retention of function. A. We transformed the strain carrying a chromosomal *mGFP-rep* allele with a derivative of pRC7 carrying a wild-type *rep* allele (pAM403), which is an unstable low copy plasmid that also carries the *lacZYA* genes (3). Presence of this plasmid confers blue colour to the colonies on Xgal indicator plates in strains chromosomally deleted for the *lac* operon. In rapidly growing cells, *rep* or *uvrD* is essential for viability, while loss of both is inviable (4, 9). Therefore, *ΔuvrD* cells with a functional *rep* allele are viable and can readily lose pRC7rep, giving rise to white colonies on LB Xgal IPTG plates, but cells lacking *rep* function cannot lose pRC7rep (4, 10) (see Ai and ii). The *mGFP-rep ΔuvrD* cells produced white colonies on Xgal media, indicating that the *mGFP-rep* fusion was functional (see Aiii). B. We also generated an *mGFP-priC* fusion and tested for retention of function by introducing a pRC7 derivative carrying the wild-type *priC* allele (pAM421) in a *ΔpriB ΔlacZYA* strain. Cells require either functional PriB or PriC for viability but loss of both is lethal (11) (See Bi and ii). *mGFP-priC ΔpriB* cells could lose pRC7priC indicating a functional PriC fusion protein (See Biii).

**Figure S2.**
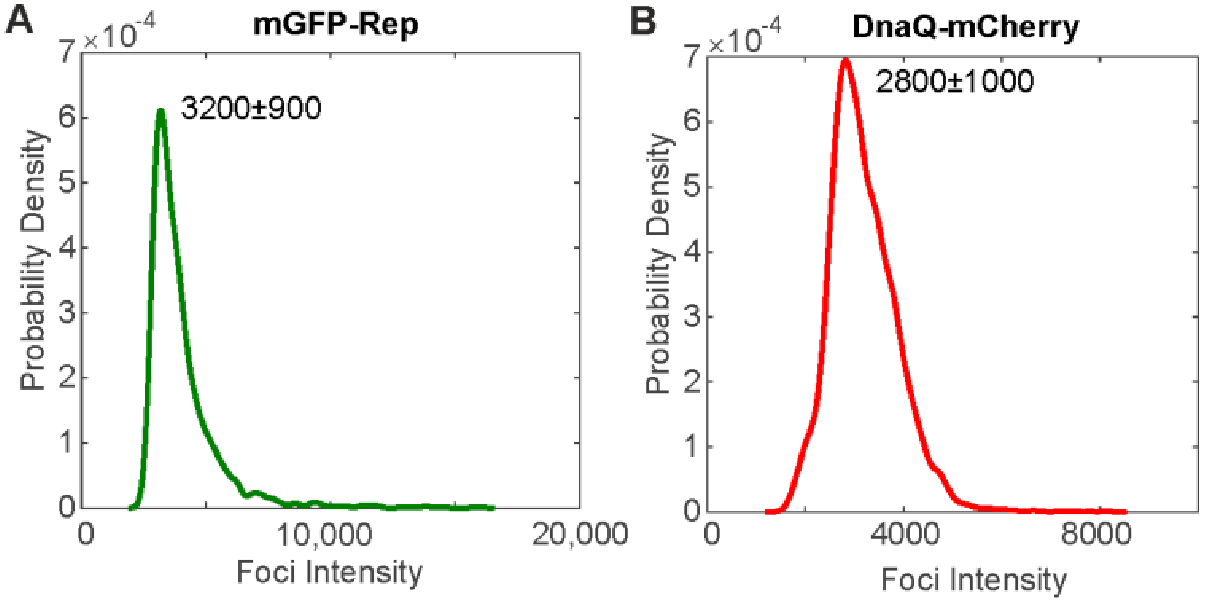
Brightness of single mGFP and mCherry molecules. A. and B. Characteristic intensity distributions rendered as kernel density estimates of single *mGFP-rep* and *mCherry-DnaQ*. Peak±full width at half maximum indicated. Distributions calculated from the tracked foci intensity distributions from the end of the photobleach process such that only single fluorophore molecules are detected. Number of molecules per foci before bleaching is determined by dividing the initial foci intensity by these values for the equivalent fluorophore.

**Figure S3.**
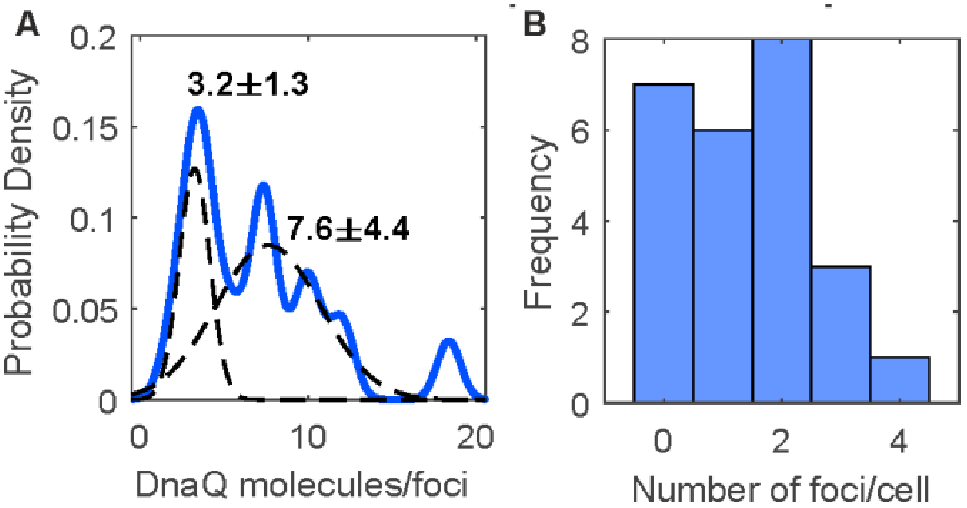
DnaQ-mGFP Slimfield analysis. A. KDE of DnaQ foci stoichiometry with double Gaussian fits in dotted lines, peak values ± SE indicated. B. Histogram showing the number of DnaQ foci detected per cell.

**Figure S4.**
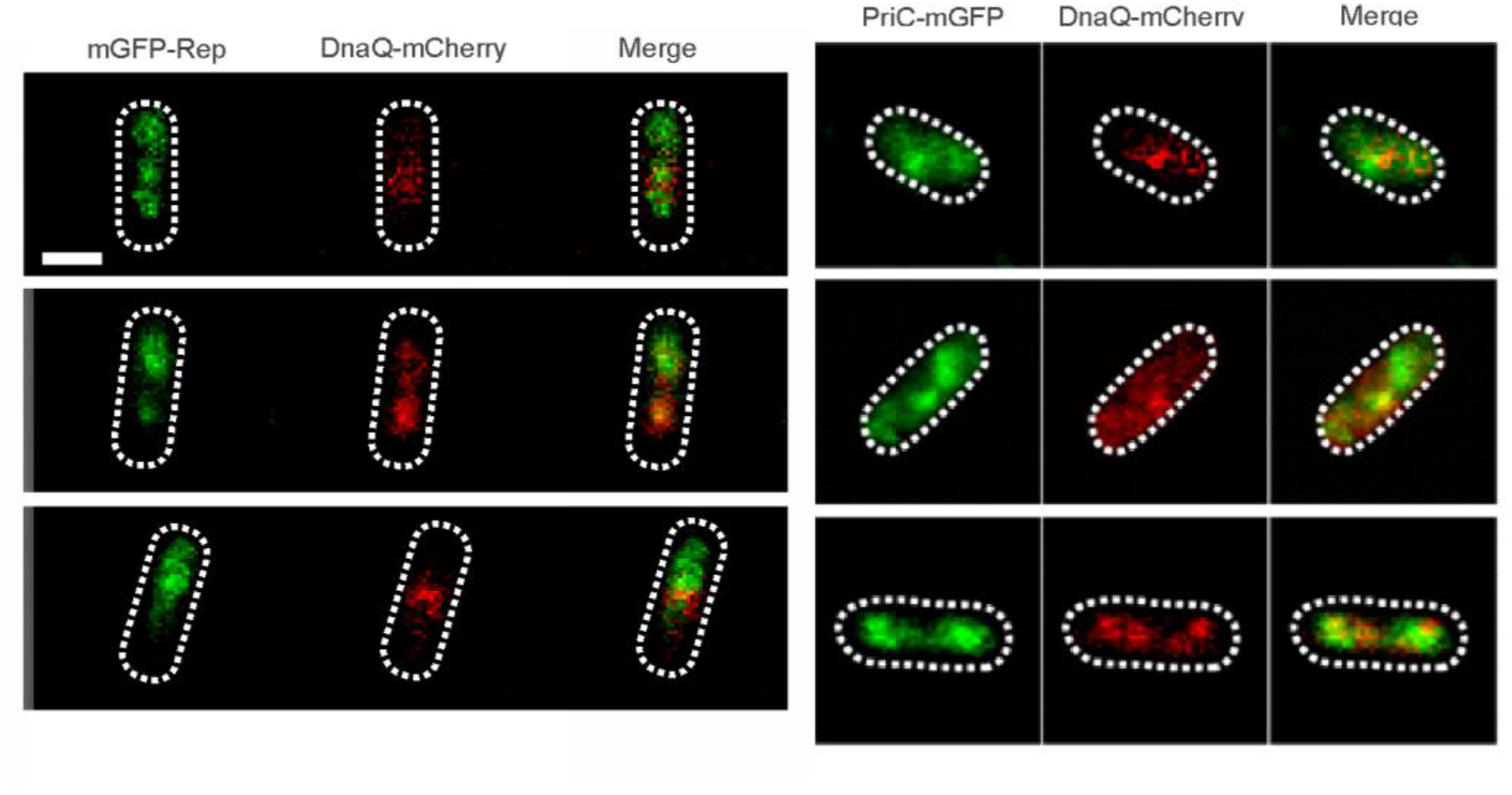
Rep and PriC localization. Dual color Slimfield images of mGFP-Rep:DnaQ-mCherry (left panel) and mGFP-PriC:DnaQ-mCherry (right panel), scale bar 1 micron.

**Figure S5.**
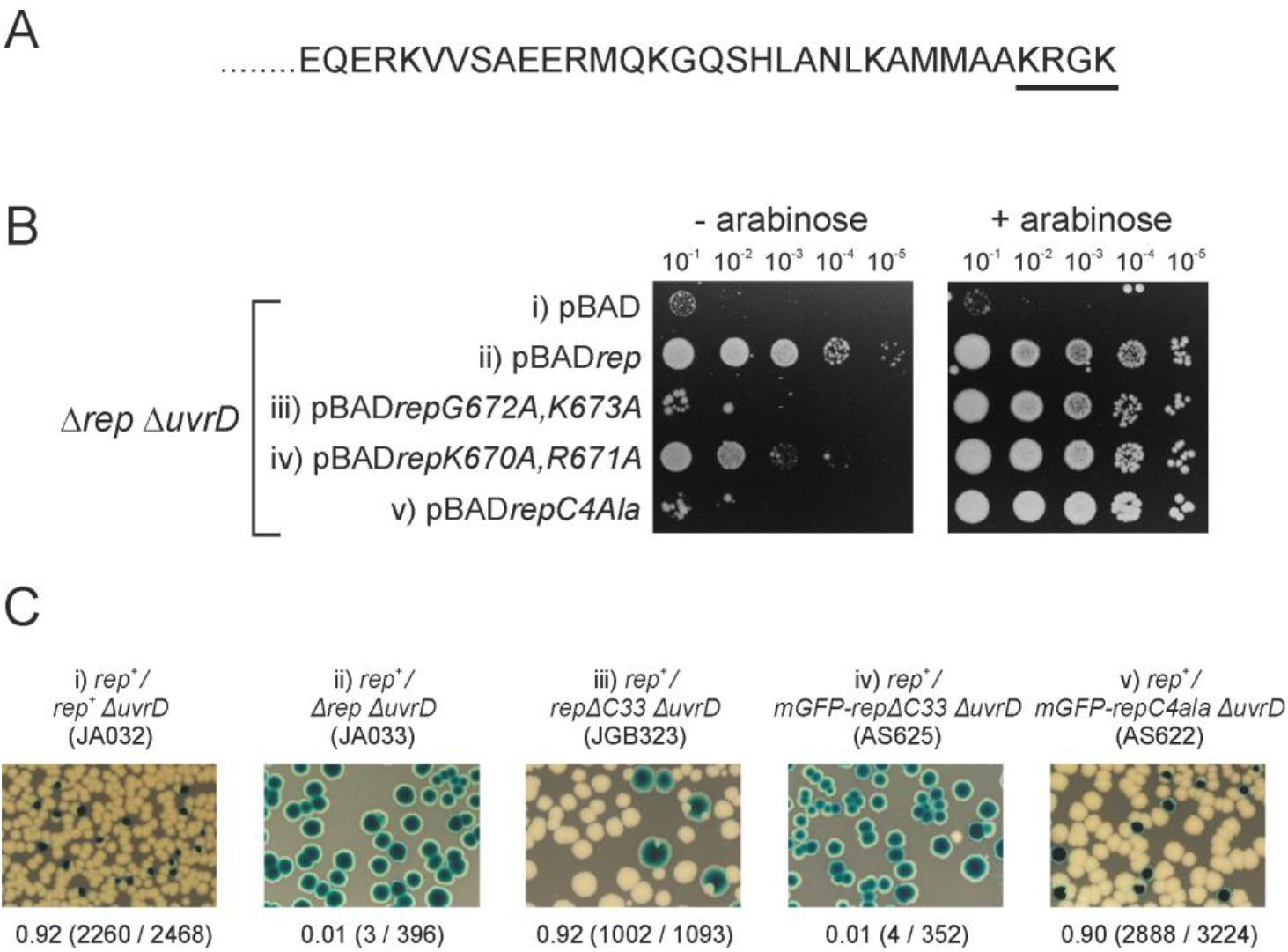
Generation of a *rep* mutant that phenocopies *repΔC33* but which does not lose function when fused to mGFP. A. The C-terminal 33 amino acids of Rep within which the DnaB interaction domain resides (4). The final four residues of this C-terminal region, underlined, were chosen as targets for mutagenesis based on their unusually high charge density. B. The plasmid pBAD bears an arabinose-inducible promoter that provides very low and very high levels of expression of genes when arabinose is absent or present in the growth medium, respectively (4). Strains lacking both the *rep* and *uvrD* genes are inviable when grown rapidly on rich medium (4, 9) and so *Δ*rep *Δ*uvrD/pBAD cells cannot form colonies on LB agar due to absence of a complementing helicase gene in pBAD (4) (see also i). In contrast, pBADrep, encoding wild type *rep*, allows *Δrep ΔuvrD* cells to grow on LB agar in both the absence and presence of arabinose, consonant with very low levels of *rep* gene expression being sufficient to sustain viability (4) (see also ii). Absence of the Rep-DnaB interaction is characterised by loss of complementation at very low levels of helicase expression (- arabinose) but maintenance of complementation at high expression levels (+ arabinose) (4). We used this pattern of complementation as a readout of the interaction between Rep and DnaB. We introduced pairs of alanine mutations into the final four codons within pBADrep. Both pBAD*repG672A,K673A* and pBAD*repK670A,R671A* displayed reduced complementation in the absence of arabinose but full complementation with arabinose, indicating involvement of both pairs of residues in the Rep-DnaB interaction (iii and iv). We therefore constructed *pBADrepC4Ala*, in which all four C-terminal residues are mutated to alanine and found that complementation required arabinose (Bv). These data indicate that the final 4 amino acids within the Rep C-terminal region are the residues that determine the phenotype displayed by *repΔC33*. C To determine whether an mGFP-repC4Ala fusion retains function, we employed a plasmid loss assay to determine the viability of strains. pRC7 is a highly unstable, very low copy plasmid that encodes *lacIZYA* (3). Retention or loss of this plasmid can be monitored by blue/white colony colour in strains bearing a chromosomal deletion of the lac operon. pAM403 is a derivative of pRC7 encoding wild type *rep* (12). *rep^+^ΔuvrD* cells can lose pRC7*rep* rapidly under rapid growth conditions, forming white colonies on LB X-gal IPTG plates, whereas *Δ*rep *ΔuvrD* cells can grow only if they retain pRC7*rep* (4, 10) (see also Ci and ii). *repΔC33 ΔuvrD* cells are viable since native expression levels of *repΔC33* are sufficient to retain partial accessory helicase function (13, 14). However, fusion of *repΔC33* to mGFP resulted in much lower viability than the original *repΔC33* allele (compare iv with iii). In contrast, *mGFP-repC4Ala ΔuvrD* cells retained viability indicating that the mGFP fusion did not have an adverse effect on RepC4Ala function (v).

**Figure S6.**
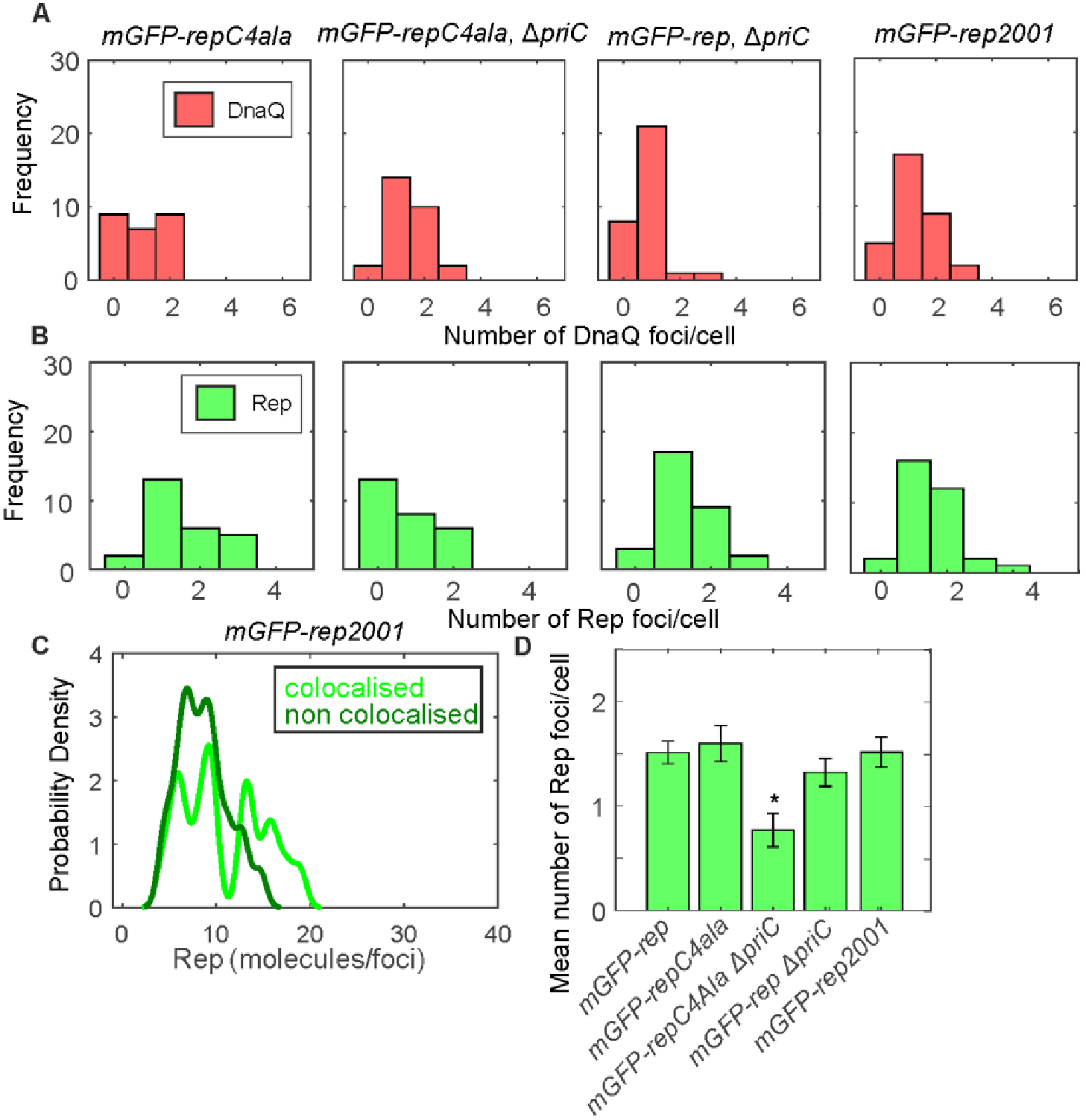
Colocalization analysis of Rep mutants. A. Number of detected DnaQ foci/cell and B. Number of detected Rep foci/cell in the absence and presence of *repC4Ala, ΔpriC* and *rep2001* C. Kernel density estimates of the number of mGFP-Rep molecules in foci colocalised with DnaQ-mCherry (light green lines) and foci not colocalized with DnaQ-mCherry (dark green lines) in mGFP-rep2001. D. The mean number of mGFP-rep foci detected per cell for wild type and mutant strains. SE indicated.

**Figure S7.**
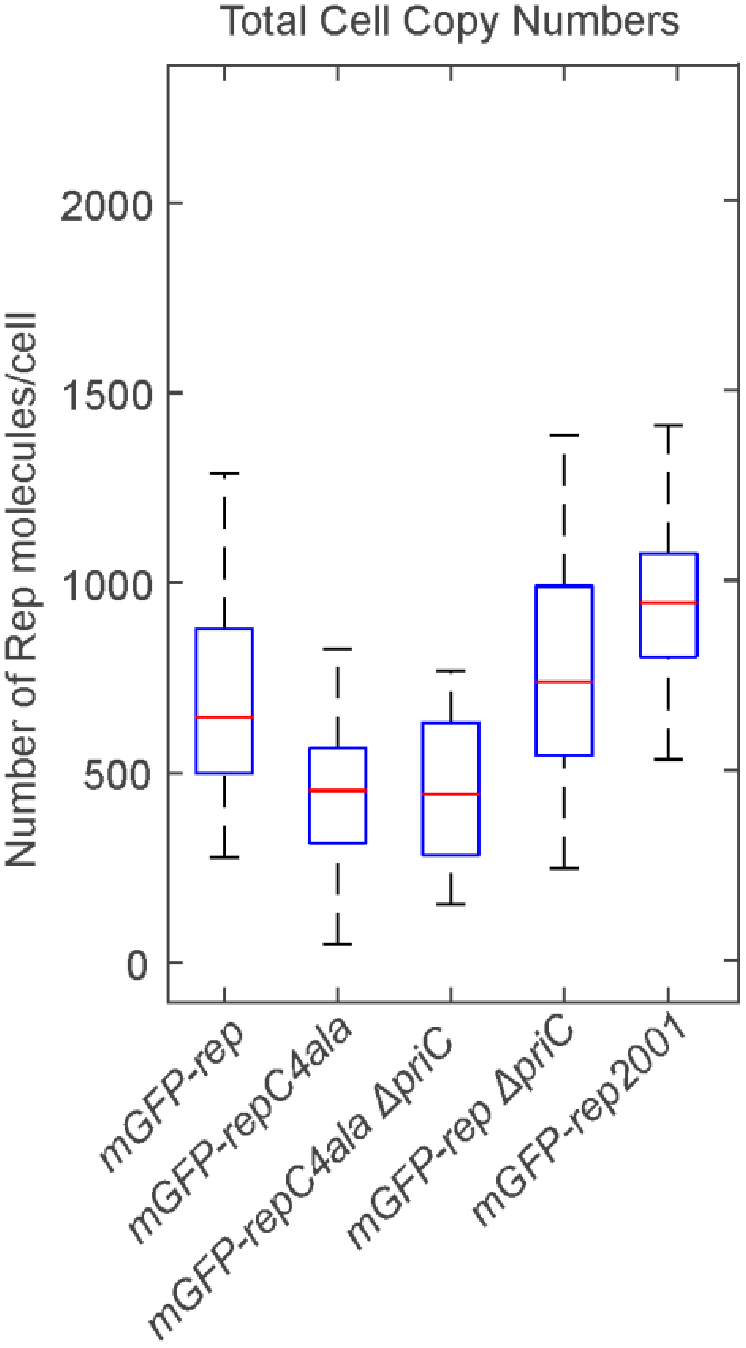
Total cell copy numbers. Boxplot of the total number of mGFP-Rep molecules per cell, estimated by numerical integration of the whole cell fluorescence. Median is shown in red, bottom and top of the blue box mark the 25^th^ and 75^th^ percentiles and whiskers extend to the most extreme points not considered outliers (2.7 standard deviations covering 99.3% of normally distributed data).

**Figure S8.**
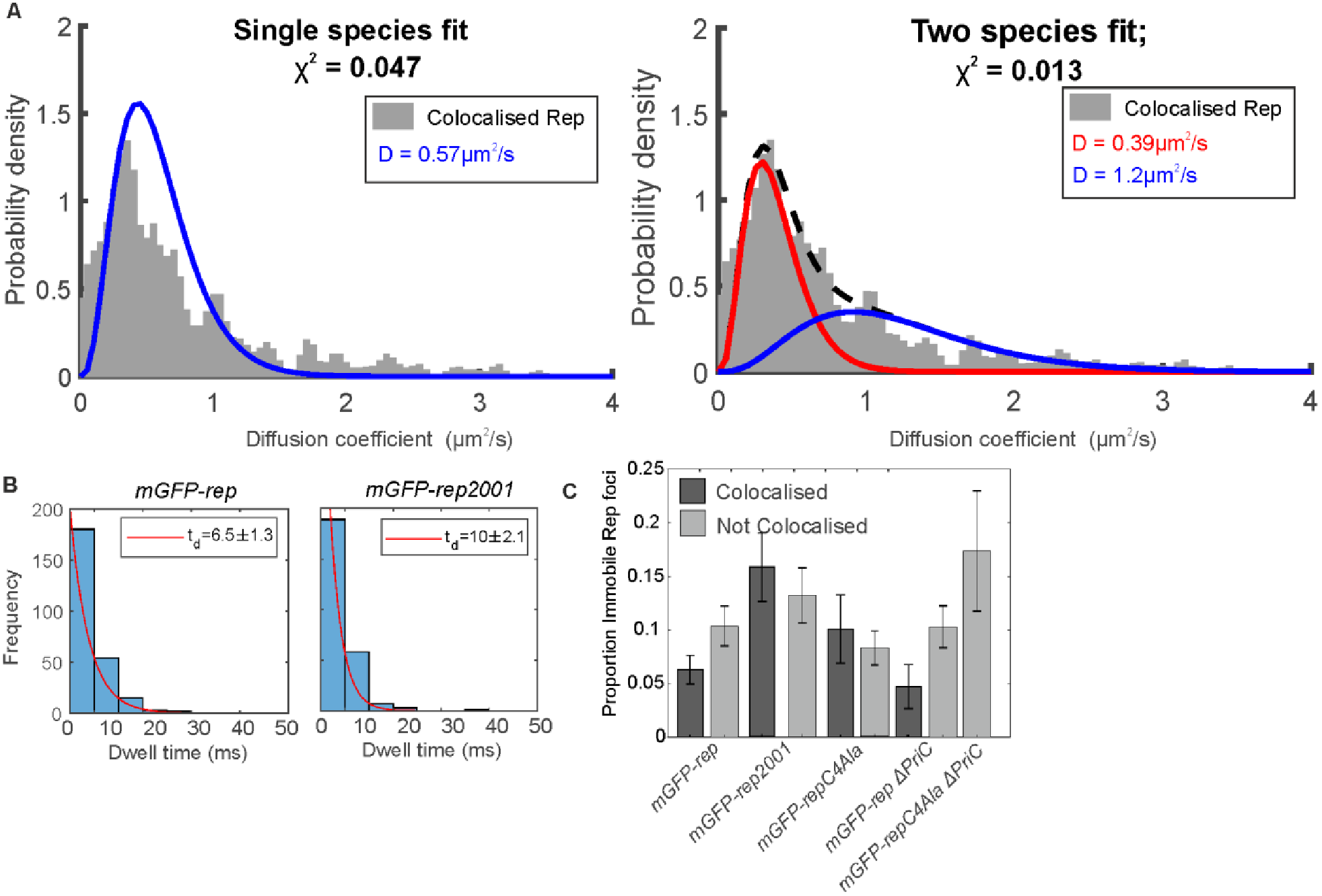
Mobility analysis of Rep mutant. A. 1 and 2 species (diffusion coefficients in this case) fits to colocalized Rep diffusion coefficients with larger reduced chi^2^ than the 3 species fit shown in Fig. 4. Thus 3 species fits were used for all the data. B. *mGFP-rep* foci dwell time with mCherry-DnaQ foci distribution with an exponential fit (red) in the absence and presence of *rep2001* C. Proportion of immobile colocalized and non colocalized mGFP-Rep in the absence and presence of *repCΔAla, ΔpriC* and *rep2001* Error bars given by the 95% confidence intervals of the fit.

### SI Appendix

#### Strain construction

All strains used in this study are derivatives of the laboratory wild-type strain TB28. Briefly, for tagging *dnaQ, linker-mGFPmut3* followed by a kanamycin resistance cassette flanked by *frt* sites was amplified by PCR from the plasmid pDHL580 using primers oAS77 and oAS79 (SI Table S4), and *linker-mCherry-<kan>* was amplified from pJGB374 using primers oAS132 and oAS133. The amplification primers had a 50 bp homology at their 5’ end to the last 50 bp of the dnaQ gene preceding the stop codon (forward primer) or the 50 bp immediately after the stop codon (reverse primer). The resulting PCR products thus had homology either side such that recombination with the chromosome would result in in-frame integration of *linker-mGFP-<kan>* and *linker-mCherry-<kan>* immediately downstream of *dnaQ*, resulting in *dnaQ-mGFP-<kan>* and *dnaQ-mCherry-<kan>* alleles.

The PCR products were treated with *Dpn*l, gel purified, and introduced by electroporation into cells expressing the lambda Red genes from the plasmid pKD46. The recombinants were selected for kanamycin resistance and screened for ampicillin sensitivity. The colonies obtained were verified for integration by PCR and sequencing with primers oAS84 and oAS85.

*mGFP-rep-<kan>* fusions for various *rep* alleles were amplified from plasmids pAS79 *(rep^+^)*, pAS124 *(repC4ala)* and pAS127 *(rep2001)* with primers oAS141 and oJGB380 having 50 bp homology on either end of the native *rep* locus. Likewise *mCherry-rep-<kan>* was amplified from pJGB380 using primers oJGB379 and oJGB380. *mGFP-priC-<kan>* was amplified from the plasmid pAS65 using primers oAS136 and oJGB389. All PCR products were introduced on the chromosome of cells expressing lambda red genes at the native loci after *Dpn*l digestion, gel extraction, and electroporation as described above for *dnaQ* fusions. The *rep* recombinants were verified by PCR amplification and sequencing using the primers oJGB418, oMKG70, oMKG71, oPM363, oPM372, and oPM376. The *priC* recombinants were verified by PCR amplification and sequencing with primers oJGB402, oJGB403, oJGB417, and oJGB418.

Where required, the kanamycin resistance gene was removed by expressing Flp recombinase from the plasmid pCP20 (7) to generate kanamycin sensitive strains carrying the FP fusions.

Dual labeled strains were created by introducing the kanamycin tagged FP alleles by standard P1 mediated transduction into single labelled strains carrying the required FP allele after removing the linked kanamycin marker.

All plasmids used in this study are listed in SI Table S3 and all primers are listed in SI Table S4.

##### RepC4Ala

pBAD is a plasmid conferring kanamycin resistance that contains an arabinose-inducible promoter upstream of a multiple cloning site whilst pBADrep is a derivative encoding wild type rep (4). *pBADrepG672A,K673A* and *pBADrepK670A,R671A* were constructed by site-directed mutagenesis of the indicated codons within pBADrep. *pBADrepC4Ala* is a derivative of pBADrep in which all four codons were altered by site-directed mutagenesis to encode alanine. Assays to determine the ability of pBAD and derivatives to complement *Δrep ΔuvrD* inviability on rich medium were performed as described (4). Plasmid loss experiments to determine the viability of combinations of chromosomal alleles were performed as described (12).

##### Determination of generation time of the *E. coli* strains by analysis of growth

Cells were grown overnight in LB medium at 37°C at 200rpm. The saturated overnight cultures were washed once with 1× 56 salts and diluted 100 fold in fresh 1× 56 salts with 0.2% glucose as the carbon source. Aliquots of 100 μl each of the diluted cultures in the fresh medium were pipetted into individual wells of a 96 well clear flat bottom sterile microplate (Corning). The microplate containing the diluted cultures was incubated in a BMG LABTECH SPECTROstar Nano microplate reader at 37°C and the optical density (A600) values were recorded at defined time intervals. The time taken for the optical density values to double during the exponential growth phase of the culture was taken as the generation time. The values expressed are the means of three independent replicates, with the standard deviation and standard errors indicated.

##### Single-molecule microscopy and analysis

A dual color bespoke single-molecule microscope was used (15) which used a narrow 10μm at full width half maximum excitation field at the sample plane to generate Slimfield illumination. Excitation was from 488nm and 561nm 50mW Obis lasers digitally modulated to produce alternating laser excitation with 5ms period. Modulation was produced by National Instruments dynamic I/O module NI 9402. Excitation was coupled into a Zeiss microscope body with a Mad City Lab’s nanostage holding the sample. Emission was magnified to 80nm/pixel and imaged using an Andor Ixon 128 emCCD camera. Green/Red images were split using a bespoke colour splitter consisting of a dual-pass green/red dichroic mirror centered at long-pass wavelength 560nm and emission filters with 25nm bandwidths centered at 542nm and 594nm.

Samples were imaged on agarose pads suffused with media as described previously (16).

Foci were automatically detected and tracked using bespoke Matlab software described previously (17). In brief bright foci were identified by image transformation and thresholding. The centroid of candidate foci were determined using iterative Gaussian masking (18) and accepted if their intensity was greater than a signal to noise ratio (SNR) of 0.4. Intensity was defined as the summed pixel intensity inside a 5 pixel circular region of interest (ROI) corrected for the background in an outer square ROI of 17×17 pixels. SNR was defined as the mean BG corrected pixel intensity in the circular ROI divided by the standard deviation in the square ROI. Foci were linked together into trajectories between frames if they were within 5 pixels of each other.

Stoichiometry was determined by fitting the first 3 intensity values of a foci to a straight line, using the intercept as the initial intensity and dividing this by the characteristic intensity of GFP or mCherry. This characteristic intensity was determined from the distribution of foci intensity values towards the end of the photobleach confirmed by overtracking foci beyond their bleaching to generate individual photobleach steps of the characteristic intensity (Fig S2).

Red and green images were aligned based on the peak of the 2D cross correlation between brightfield images. Colocalization between foci and the probability of random colocalization was determined as described previously (19).

Microscopic diffusion coefficients were calculated by fitting the first 3 mean square displacement (MSD) values, i.e. equivalent to time interval values of 5, 10 and 15 ms, with a linear fit constrained through the equivalent localization precision MSD (20).

The upper bound of stoichiometry in the pool was calculated using an approach modified from previously (15). We modeled an average *E.coli* cell volume as equivalent to a cylinder of diameter 1 μm and length which varies between 1–4 μm depending on the stage in the cell cycle, capped by 2 hemispheres (21). This morphology indicates a mean volume *V* of 3.7–13.1 μm^3^ per cell, assumed largely accessible to Rep unlike far large protein complexes such as polysomes which exhibit nucleoid exclusion (22). If the mean cell copy number for in the pool for Rep is *n* with mean foci stoichiometry of *S* then the mean number of Rep foci *F* in the pool is *n/S*. If each Rep focus occupies an equivalent sphere of radius r such that the sum of all spheres is equivalent to the cell volume then F.4πr^3^/3=V. The optical resolution limit, identified as the pointed spread function width *w* of our microscope, for our setup was measured previously for mGFP excitation to be ~230 nm (15). For Rep foci to be part of the pool implies that the mean nearest neighbour foci separation (i.e. 2r) is not greater than w, such that *r* is the radius of the sphere associated with each focus with the sum of all such spheres having a total volume V. Thus, assuming relative insensitivity to blur artefacts(3) with our rapid sampling:

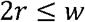

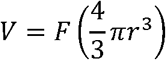

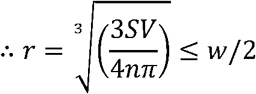

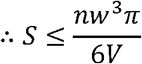

Using an average value of *n* of ~650 molecules per cell (SI Fig. 7) the range in *V* suggests an upper limit to *S* in the range 0.3–1.1 molecules per Rep focus, consistent with a monomeric pool for Rep, to the nearest integer.

#### Supplementary Movie legends

**Supplementary Movies 1 and 2**. Slimfield movies indicating Rep (green) and DnaQ (red) dynamic localization on two separate *E. coli* cells. Red channel data has been removed after the first five image frames, with a white circle indicating the position of a detect DnaQ focus, marking the location of a replication fork.

Author contributions
P.M. and M.C.L. designed research; A.S., A.J.M.W, A.H. and J.-G.B performed research; A.J.M.W and A.S.; A.S., A.J.M.W., P.M. and M.C.L. wrote the paper.

## References

1. McGlynn P, Savery NJ, & Dillingham MS (2012) The conflict between DNA replication and transcription. Mol. Microbiol. 85(1):12–20.

2. Hamperl S & Cimprich KA (2016) Conflict Resolution in the Genome: How Transcription and Replication Make It Work. Cell 167(6):1455–1467.

3. Yeeles JT, Poli J, Marians KJ, & Pasero P (2013) Rescuing stalled or damaged replication forks. Cold Spring Harb. Perspect. Biol. 5(5):a012815.

4. Bruning JG, Howard JL, & McGlynn P (2014) Accessory Replicative Helicases and the Replication of Protein-Bound DNA. J. Mol. Biol. 426(24):3917–3928.

5. Ivessa AS, et al. (2003) The *Saccharomyces cerevisiae* helicase Rrm3p facilitates replication past nonhistone protein-DNA complexes. Mol. Cell 12(6):1525–1536.

6. Gupta MK, et al. (2013) Protein-DNA complexes are the primary sources of replication fork pausing in *Escherichia coli*. Proc. Natl. Acad. Sci. U S A 110(18):7252–7257.

7. Azvolinsky A, Giresi PG, Lieb JD, & Zakian VA (2009) Highly transcribed RNA polymerase II genes are impediments to replication fork progression in *Saccharomyces cerevisiae*. Mol. Cell 34(6):722–734.

8. Ivessa AS, Zhou JQ, Schulz VP, Monson EK, & Zakian VA (2002) *Saccharomyces* Rrm3p, a 5’ to 3’ DNA helicase that promotes replication fork progression through telomeric and subtelomeric DNA. Genes Dev. 16(11):1383–1396.

9. Guy CP, et al. (2009) Rep Provides a Second Motor at the Replisome to Promote Duplication of Protein-Bound DNA. Mol. Cell 36(4):654–666.

10. Boubakri H, de Septenville AL, Viguera E, & Michel B (2010) The helicases DinG, Rep and UvrD cooperate to promote replication across transcription units *in vivo*. EMBO J. 29(145–157).

11. Chodavarapu S & Kaguni JM (2016) Replication Initiation in Bacteria. Enzymes 39: 1–30.

12. Gilhooly NS, Gwynn EJ, & Dillingham MS (2013) Superfamily 1 helicases. Front. Biosci. (Schol. Ed.) 5: 206–216.

13. Bruning JG, Howard JAL, Myka KK, Dillingham MS, & McGlynn P (2018) The 2B subdomain of Rep helicase links translocation along DNA with protein displacement. Nucleic acids research.

14. Yarranton GT & Gefter ML (1979) Enzyme-catalyzed DNA unwinding: studies on *Escherichia coli* rep protein. Proc. Natl. Acad. Sci. U S A 76(4):1658–1662.

15. LeBowitz JH & McMacken R (1986) The *Escherichia coli* dnaB replication protein is a DNA helicase. J. Biol. Chem. 261(10):4738–4748.

16. Schmidt KH & Kolodner RD (2004) Requirement of Rrm3 helicase for repair of spontaneous DNA lesions in cells lacking Srs2 or Sgs1 helicase. Mol. Cell. Biol. 24(8):3213–3226.

17. Torres JZ, Schnakenberg SL, & Zakian VA (2004) *Saccharomyces cerevisiae* Rrm3p DNA helicase promotes genome integrity by preventing replication fork stalling: viability of rrm3 cells requires the intra-S-phase checkpoint and fork restart activities. Mol. Cell. Biol. 24(8):3198–3212.

18. Uzest M, Ehrlich SD, & Michel B (1995) Lethality of *rep recB* and *rep recC* double mutants of *Escherichia coli*. Mol. Microbiol. 17(6):1177–1188.

19. Byrd AK & Raney KD (2004) Protein displacement by an assembly of helicase molecules aligned along single-stranded DNA. Nat. Struct. Mol. Biol. 11(6):531–538.

20. Sabouri N, McDonald KR, Webb CJ, Cristea IM, & Zakian VA (2012) DNA replication through hard-to-replicate sites, including both highly transcribed RNA Pol II and Pol III genes, requires the *S. pombe* Pfh1 helicase. Genes Dev. 26(6):581–593.

21. Steinacher R, Osman F, Dalgaard JZ, Lorenz A, & Whitby MC (2012) The DNA helicase Pfh1 promotes fork merging at replication termination sites to ensure genome stability. Genes Dev. 26(6):594–602.

22. Schmidt KH, Derry KL, & Kolodner RD (2002) *Saccharomyces cerevisiae* RRM3, a 5’ to 3’ DNA helicase, physically interacts with proliferating cell nuclear antigen. J. Biol. Chem. 277(47):45331–45337.

23. Azvolinsky A, Dunaway S, Torres JZ, Bessler JB, & Zakian VA (2006) The *S. cerevisiae* Rrm3p DNA helicase moves with the replication fork and affects replication of all yeast chromosomes. Genes Dev. 20(22):3104–3116.

24. McDonald KR, et al. (2016) Pfh1 Is an Accessory Replicative Helicase that Interacts with the Replisome to Facilitate Fork Progression and Preserve Genome Integrity. PLoS Genet. 12(9):e1006238.

25. Atkinson J, Gupta MK, & McGlynn P (2011) Interaction of Rep and DnaB on DNA. Nucleic Acids Res. 39(4):1351–1359.

26. Bruning JG, Howard JA, & McGlynn P (2016) Use of streptavidin bound to biotinylated DNA structures as model substrates for analysis of nucleoprotein complex disruption by helicases. Methods 108: 48–55.

27. Johnson A & O’Donnell M (2005) Cellular DNA replicases: components and dynamics at the replication fork. Annu. Rev. Biochem. 74: 283–315.

28. Bentchikou E, et al. (2015) Are the SSB-Interacting Proteins RecO, RecG, PriA and the DnaB-Interacting Protein Rep Bound to Progressing Replication Forks in *Escherichia coli*? PloS one 10(8):e0134892.

29. Sandler SJ (2000) Multiple genetic pathways for restarting DNA replication forks in *Escherichia coli* K-12. Genetics 155(2):487–497.

30. Heller RC & Marians KJ (2005) Unwinding of the nascent lagging strand by Rep and PriA enables the direct restart of stalled replication forks. J. Biol. Chem. 280(40):34143–34151.

31. Windgassen TA, Wessel SR, Bhattacharyya B, & Keck JL (2018) Mechanisms of bacterial DNA replication restart. Nucleic acids research 46(2):504–519.

32. Datsenko KA & Wanner BL (2000) One-step inactivation of chromosomal genes in *Escherichia coli* K-12 using PCR products. Proc. Natl. Acad. Sci. U S A 97(12):6640–6645.

33. Zacharias DA, Violin JD, Newton AC, & Tsien RY (2002) Partitioning of lipid-modified monomeric GFPs into membrane microdomains of live cells. Science 296(5569):913–916.

34. Plank M, Wadhams GH, & Leake MC (2009) Millisecond timescale slimfield imaging and automated quantification of single fluorescent protein molecules for use in probing complex biological processes. Integr. Biol. (Camb.) 1(10):602–612.

35. Miller H, Zhou Z, Shepherd J, Wollman A, & Leake M (2017) Single-molecule techniques in biophysics: a review of the progress in methods and applications. Reports on progress in physics. Physical Society.

36. Reyes-Lamothe R, Sherratt DJ, & Leake MC (2010) Stoichiometry and architecture of active DNA replication machinery in *Escherichia coli*. Science 328(5977):498–501.

37. Badrinarayanan A, Reyes-Lamothe R, Uphoff S, Leake MC, & Sherratt DJ (2012) *In vivo* architecture and action of bacterial structural maintenance of chromosome proteins. Science 338(6106):528–531.

38. Lund VA, et al. (2018) Molecular coordination of Staphylococcus aureus cell division. eLife 7.

39. Wollman AJ, et al. (2017) Transcription factor clusters regulate genes in eukaryotic cells. eLife 6.

40. Miller H, et al. (2018) High-Speed Single-Molecule Tracking of CXCL13 in the B-Follicle. Frontiers in immunology 9: 1073.

41. Reyes-Lamothe R, Possoz C, Danilova O, & Sherratt DJ (2008) Independent positioning and action of *Escherichia coli* replisomes in live cells. Cell 133(1):90–102.

42. McInerney P, Johnson A, Katz F, & O’Donnell M (2007) Characterization of a triple DNA polymerase replisome. Mol. Cell 27(4):527–538.

43. Georgescu RE, Kurth I, & O’Donnell ME (2011) Single-molecule studies reveal the function of a third polymerase in the replisome. Nature Structural & Molecular Biology 19(1):113–116.

44. Llorente-Garcia I, et al. (2014) Single-molecule *in vivo* imaging of bacterial respiratory complexes indicates delocalized oxidative phosphorylation. Biochim. Biophys. Acta 1837(6):811–824.

45. Wollman AJ & Leake MC (2015) Millisecond single-molecule localization microscopy combined with convolution analysis and automated image segmentation to determine protein concentrations in complexly structured, functional cells, one cell at a time. Faraday discussions 184: 401–424.

46. Atkinson J, et al. (2011) Localization of an accessory helicase at the replisome is critical in sustaining efficient genome duplication. Nucleic Acids Res. 39(3):949–957.

47. Singleton MR, Dillingham MS, & Wigley DB (2007) Structure and mechanism of helicases and nucleic acid translocases. Annu. Rev. Biochem. 76: 23–50.

48. Ludlam AV, McNatt MW, Carr KM, & Kaguni JM (2001) Essential Amino Acids of *Escherichia coli* DnaC Protein in an N-terminal Domain Interact with DnaB Helicase. J. Biol. Chem. 276(29):27345–27353.

49. Marszalek J & Kaguni JM (1994) DnaA protein directs the binding of DnaB protein in initiation of DNA replication in *Escherichia coli*. J. Biol. Chem. 269(7):4883–4890.

50. Wahle E, Lasken RS, & Kornberg A (1989) The dnaB-dnaC replication protein complex of *Escherichia coli*. II. Role of the complex in mobilizing dnaB functions. J. Biol. Chem. 264(5):2469–2475.

51. Wahle E, Lasken RS, & Kornberg A (1989) The dnaB-dnaC replication protein complex of *Escherichia coli*. I. Formation and properties. J. Biol. Chem. 264(5):2463–2468.

52. Arias-Palomo E, O’Shea VL, Hood IV, & Berger JM (2013) The Bacterial DnaC Helicase Loader Is a DnaB Ring Breaker. Cell 153(2):438–448.

53. Beattie TR, et al. (2017) Frequent exchange of the DNA polymerase during bacterial chromosome replication. eLife 6.

54. Brendza KM, et al. (2005) Autoinhibition of *Escherichia coli* Rep monomer helicase activity by its 2B subdomain. Proc. Natl. Acad. Sci. U S A 102(29):10076–10081.

55. Aramaki T, et al. (2013) Domain separation and characterization of PriC, a replication restart primosome factor in *Escherichia coli*. Genes Cells 18(9):723–732.

56. Wessel SR, et al. (2013) PriC-mediated DNA replication restart requires PriC complex formation with the single-stranded DNA-binding protein. J. Biol. Chem. 288(24):17569–17578.

57. Wessel SR, et al. (2016) Structure and Function of the PriC DNA Replication Restart Protein. J. Biol. Chem. 291(35):18384–18396.

58. Leake MC, Wilson D, Bullard B, & Simmons RM (2003) The elasticity of single kettin molecules using a two-bead laser-tweezers assay. FEBS letters 535(1–3):55–60.

## Supplementary References

1. Baba T, et al. (2006) Construction of *Escherichia coli* K-12 in-frame, single-gene knockout mutants: the Keio collection. Mol. Syst. Biol. 2:2006 0008.

2. Bachmann BJ (1996) Derivations and genotypes of some mutant derivatives of *Escherichia coli* K-12. Escherichia coli and Salmonella cellular and molecular biology, eds Neidhardt FC, Curtiss III R, Ingraham JL, Lin ECC, Low KB, Magasanik B, Reznikoff WS, Riley M, Schaechter M, & Umbarger HE (ASM Press, Washington, DC), Second Ed, pp 2460–2488.

3. Bernhardt TG & de Boer PA (2004) Screening for synthetic lethal mutants in Escherichia coli and identification of EnvC (YibP) as a periplasmic septal ring factor with murein hydrolase activity. Molecular microbiology 52(5):1255–1269.

4. Guy CP, et al. (2009) Rep Provides a Second Motor at the Replisome to Promote Duplication of Protein-Bound DNA. Mol. Cell 36(4):654–666.

5. Mahdi AA, Briggs GS, & Lloyd RG (2012) Modulation of DNA damage tolerance in *Escherichia coli recG* and *ruv* strains by mutations affecting PriB, the ribosome and RNA polymerase. Mol. Microbiol. 86(3):675–691.

6. Rudolph CJ, Upton AL, Stockum A, Nieduszynski CA, & Lloyd RG (2013) Avoiding chromosome pathology when replication forks collide. Nature 500(7464):608–611.

7. Datsenko KA & Wanner BL (2000) One-step inactivation of chromosomal genes in *Escherichia coli* K-12 using PCR products. Proc. Natl. Acad. Sci. U S A 97(12):6640–6645.

8. Landgraf D, Okumus B, Chien P, Baker TA, & Paulsson J (2012) Segregation of molecules at cell division reveals native protein localization. Nat. Methods 9(5):480–482.

9. Boubakri H, de Septenville AL, Viguera E, & Michel B (2010) The helicases DinG, Rep and UvrD cooperate to promote replication across transcription units *in vivo*. EMBO J. 29(145–157).

10. Myka KK, et al. (2017) Inhibiting translation elongation can aid genome duplication in *Escherichia coli*. Nucleic Acids Res. 45(5):2571–2584.

11. Sandler SJ, et al. (1999) *dnaC* mutations suppress defects in DNA replication- and recombination-associated functions in *priB* and *priC* double mutants in *Escherichia coli* K-12. Mol. Microbiol. 34(1):91–101.

12. Mahdi AA, Buckman C, Harris L, & Lloyd RG (2006) Rep and PriA helicase activities prevent RecA from provoking unnecessary recombination during replication fork repair. Genes Dev. 20(15):2135–2147.

13. Atkinson J, et al. (2011) Localization of an accessory helicase at the replisome is critical in sustaining efficient genome duplication. Nucleic Acids Res. 39(3):949–957.

14. Atkinson J, Gupta MK, & McGlynn P (2011) Interaction of Rep and DnaB on DNA. Nucleic Acids Res. 39(4):1351–1359.

15. Wollman AJ, et al. (2017) Transcription factor clusters regulate genes in eukaryotic cells. eLife 6.

16. Wollman AJ, Syeda AH, McGlynn P, & Leake MC (2016) Single-Molecule Observation of DNA Replication Repair Pathways in *E. coli*. Adv. Exp. Med. Biol. 915: 5–16.

17. Miller H, Zhou Z, Wollman AJ, & Leake MC (2015) Superresolution imaging of single DNA molecules using stochastic photoblinking of minor groove and intercalating dyes. Methods 88: 81–88.

18. Thompson RE, Larson DR, & Webb WW (2002) Precise nanometer localization analysis for individual fluorescent probes. Biophysical journal 82(5):2775–2783.

19. Llorente-Garcia I, et al. (2014) Single-molecule *in vivo* imaging of bacterial respiratory complexes indicates delocalized oxidative phosphorylation. Biochim. Biophys. Acta 1837(6):811–824.

20. Wollman AJ & Leake MC (2015) Millisecond single-molecule localization microscopy combined with convolution analysis and automated image segmentation to determine protein concentrations in complexly structured, functional cells, one cell at a time. Faraday discussions 184: 401–424.

21. Leake MC, et al. (2006) Stoichiometry and turnover in single, functioning membrane protein complexes. Nature 443(7109):355–358.

22. Bakshi S, Siryaporn A, Goulian M, & Weisshaar JC (2012) Superresolution imaging of ribosomes and RNA polymerase in live *Escherichia coli* cells. Mol. Microbiol. 85(1):21–38.

